# Looking for lipases and lipolytic organisms in low-temperature anaerobic reactors treating domestic wastewater

**DOI:** 10.1101/2021.11.16.468786

**Authors:** Reihaneh Bashiri, Ben Allen, Burhan Shamurad, Martin Pabst, Thomas P. Curtis, Irina D. Ofiţeru

**Affiliations:** School of Engineering, Newcastle University, Newcastle-upon-Tyne, NE1 7RU, UK; Department of Biotechnology, Delft University of Technology, Van der Maasweg 9, 2629HZ Delft, The Netherlands

**Keywords:** Anaerobic treatment, domestic wastewater, psychrophilic extracellular lipases, metagenomics, metaproteomics

## Abstract

Poor lipid degradation limits low-temperature anaerobic treatment of domestic wastewater even when psychrophiles are used. We combined metagenomics and metaproteomics to find lipolytic bacteria and their potential, and actual, cold-adapted extracellular lipases in anaerobic membrane bioreactors treating domestic wastewater at 4°C and 15°C. Of the 40 recovered putative lipolytic metagenome-assembled genomes (MAGs), only three (*Chlorobium, Desulfobacter*, and *Mycolicibacterium*) were common and abundant (relative abundance ≥ 1%) in all reactors. Notably, some MAGs that represented aerobic autotrophs contained lipases. Therefore, we hypothesised that the lipases we found are not always associated with exogenous lipid degradation and can have other roles such as polyhydroxyalkanoates (PHA) accumulation/degradation and interference with the outer membranes of other bacteria. Metaproteomics did not provide sufficient proteome coverage for relatively lower abundant proteins such as lipases though the expression of *fadL* genes, long-chain fatty acid transporters, was confirmed for four genera (*Dechloromonas, Azoarcus, Aeromonas* and *Sulfurimonas*), none of which were recovered as putative lipolytic MAGs. Metaproteomics also confirmed the presence of 15 relatively abundant (≥1%) genera in all reactors, of which at least 6 can potentially accumulate lipid/polyhydroxyalkanoates. For most putative lipolytic MAGs, there was no statistically significant correlation between the read abundance and reactor conditions such as temperature, phase (biofilm and bulk liquid), and feed type (treated by ultraviolet light or not). Results obtained by metagenomics and metaproteomics did not confirm each other and further work is required to identify the true lipid degraders in these systems.

## 1. Introduction

Anaerobic treatment of domestic wastewater generates energy and produces much less sludge than the conventional activated sludge process. However, it is still rarely adopted at full-scale outside of South America (Aquino et al. 2019) where the reactors work well at ambient temperatures (23°C-34°C). Therefore, for 60% of the world population that lives in countries with a temperate climate, using anaerobic treatment for treating domestic wastewater is problematic.

The major limitation is related to the first step in anaerobic digestion: hydrolysis. In this step, fermentative bacteria degrade large biopolymers like carbohydrates, proteins, and lipids, presumably by producing extracellular enzymes. Yet, at low temperatures, the rate of biological reactions drops, and hydrolysis becomes rate-limiting. Lipids degrade more slowly than carbohydrates or proteins at low temperatures, forming the bulk of the chemical oxygen demand (COD) in the effluents (Petropoulos et al. 2018) and a lipid rich scum layer in the reactors (Soares et al. 2019). Operational interventions, such as a skimmer to remove the scum layer (Lettinga et al. 1984), may help but do not address the fundamental issues of compliance or energy recovery. The estimated methane yield from 1 g glycerol trioleate (an abundant natural lipid) is 1.08 L (at standard temperature and pressure) while for 1 g glucose this is only 0.37 L (Kim and Shin 2010).

At bench/pilot scale it is possible to treat domestic wastewater anaerobically at ambient temperatures and as low as 4 °C (Kong et al. 2021, Lim et al. 2019, Maleki et al. 2019, McAteer et al. 2020, Petropoulos et al. 2017, Ribera-Pi et al. 2020, Yang et al. 2020). However, lipids remained undegraded (unlike carbohydrates and proteins) even when the psychrophilic microbial community is used to treat domestic wastewater anaerobically at 4, 8 and 15 °C (Petropoulos et al. 2018).

A lipolytic organism must be able to produce extracellular lipases to degrade lipids, and then transport the long-chain fatty acids back into the cell to be metabolised (for example, by the *beta-oxidation* pathway). Due to the difference of the cell wall structure in Gram-negative and Gram-positive bacteria, the export of lipases and import of long-chain fatty acids may differ. For instance, in Gram-negative bacteria, long-chain fatty acid transporter proteins (*fadL*) have been identified for carrying the long-chain fatty acids to the cells. However, such transporters have not yet been identified for the Gram-positive bacteria (Salvador López and Van Bogaert 2021) but presumably exist.

Poor lipid degradation at low temperatures could be due to a lack of lipase production, a lack of lipase activity, low rates of fatty acid transport or a slow *beta-oxidation* pathway. Understanding these limitations would be easier if we knew the identity of the lipolytic bacteria and, ideally, any accompanying cheaters (bacteria that consume long-chain fatty acids without producing lipases). The present study, therefore, aims to investigate the lipolytic potential of the psychrophilic bacteria adapted to treat domestic wastewater anaerobically at 4, and 15 °C. We combined molecular biology tools such as metagenomics and metaproteomics to find potential and actual expressed lipase genes or other lipolytic biomarkers like *fadL* genes as well as lipase producers and cheaters.

## 2. Material and Methods

### 2.1. Reactor set-up

The reactor set-up, inoculation, feeding and wastewater characterization is described in details by Petropoulos et al. (2017). Four anaerobic membrane bioreactors (AnMBRs) with 1 L working volume (and their duplicates) were operated at 4 °C and 15 °C under the Sterile (treated with the ultraviolet light to exclude mesophilic biomass of the feed) and Non-sterile conditions. The reactors were inoculated by psychrophilic biomass collected from the sediment and soils of Lake Geneva, Switzerland (annual temperature range −11 – 21 °C) and Svalbard, Norway (annual temperature range −16 – 6 °C), respectively. The feed of the reactors was primary influent collected from an activated sludge plant (Tudhoe Mill, County Durham, UK).

### 2.2. Metagenomics and data analysis

DNA was extracted from the AnMBRs samples (both from the bulk liquid and biofilm formed on the membrane) using the CTAB method (Griffiths et al. 2000) and sent for sequencing (HiSeq 2500 platform) to the Earlham Institute, Norwich. Amplification free, Illumina compatible libraries were constructed using the Kapa Hyper Prep kit. Aliquots of each samples were run on two lanes/two flowcells to generate paired end (PE 250) reads of about 300 Mb.

*FastQC v0.11.5* was employed to check the quality of reads, and *Cutadapt v1.18* and *Trimmomatic v0.36* were used to trim the adapters and poor regions. Removed parts from the *Read 1* were AGATCGGAAGAGCACACGTCTGAACTCCAGTCA and from the *Read 2* were AGATCGGAAGAGCGTCGTGTAGGGAAAGAGTGT, respectively. Filtered reads were co-assembled with *MEGAHIT v.1.2.9* using a high-performance computer at Newcastle University. Obtained contigs were then binned with *MetaBat2 v1.7* to recover the metagenome-assembled genomes (MAGs). To evaluate the quality of the bins, *CheckM v1.0.18* was used and MAGs with more than 90% completeness and less than 10% contamination were selected as putative lipolytic bins. The *FASTA* file of the selected MAGs were uploaded to *KBase* (Arkin et al. 2018) and annotated using *Prokka v1.12.* After annotation, lipase genes were searched with their EC number, 3.1.1.3. MAGs that had at least one (putative) lipase gene were specified as putative lipolytic bins. Specific EC numbers of other hydrolytic enzymes like phosphatases, proteases, esterases and carbohydrate degraders were also searched (Supplementary File 1, Table S1). Further analysis like the taxonomic classification was performed by *GTDB-Tk v0.3.2* on the putative lipolytic MAGs at *KBase.* To find the relative abundance of the microorganisms existing at each reactor condition, reads from both biofilm and liquid phase of the replicate reactors at each temperature and treatment set-up were merged with *KBase* apps and were analysed with *GOTTCHA2 v2.1.5.* All statistical analysis was performed using *Minitab 18.* Metagenomics data are accessible at European Nucleotide Archive (ENA) under the accession number PRJEB47041.

### 2.3. Metaproteomics and data analysis

Before protein extraction, volatile suspended solid (VSS) was measured following the standard method of American Public Health Association. Proteins were extracted from both biofilm and bulk liquid of AnMBRs using the protocol suggested for the extraction of extracellular lipases by both Gessesse et al. (2003) and Frølund et al. (1996). The extracted proteins were quantified by Pierce™ Modified Lowry Protein Assay Kit (Thermo Fisher Scientific) prior to precipitation by the phenol/chloroform method (Wessel and Flügge 1984). Precipitated proteins were solubilized and reduced in Laemmli buffer and β-mercaptoethanol, sonicated (20 min, cool temperature) and heated (5 min, 60 °C) before being run on one-dimensional sodium dodecyl sulphate polyacrylamide gel electrophoresis, for 5 min at 120 V (Bio-Rad Mini-PROTEAN®). The gel was stained following the protocol of Bio-Safe Coomassie Brilliant Blue G-250 and was destained overnight (details in Supplementary file 2). In-gel digestion and mass spectrometry were done at Newcastle University Protein & Proteome Analysis Centre using Ab-Sciex TripleTOF 6600 mass spectrometer following the protocol detailed in the Supplementary File 2.

Mass spectrometric raw data were converted to .mgf files using *MSConvert* and analysed as a single group using *PEAKS Studio X* using a High-Performance Computing Windows workstation. The metagenomics protein sequence database was cleaned for sequence redundancy and annotation errors using *CD-hit* and *notepad*+ +. Furthermore, the database search using the cleaned metagenomics constructed database was performed using a two-round search strategy. The initial search allowed 50 ppm parent ion and 0.1 Da fragment mass error tolerance and carbamidomethylation as fixed modification. Protein matches of the initial search with a −10lgP protein score greater or equal 20 were collected, which resulted in a preliminary search output of 11,814 protein groups. The second-round search, using the refined database from the first-round search, allowed up to 3 missed cleavages, 50 ppm parent ion and 0.1 Da fragment mass error tolerance, carbamidomethylation as fixed modification, oxidation and deamidation as variable modifications and employed a decoy fusion database for determining false discovery rates. Peptide spectrum matches were filtered against 1% or 5% false discovery rates (FDR), and protein identifications with 2 or more unique peptides across the group were considered as significant matches. Processing of metadata was done using MATLAB 2017b. Additional taxonomic and KEGG number annotations was performed using *GhostKOALA v2.2.* The mass spectrometry proteomics data have been deposited to the ProteomeXchange Consortium via the PRIDE partner repository with the dataset identifier PXD028388 (Reviewer account: Username: Password: Tl8JGpJd).

## 3. Results and Discussion

### 3.1. Reads, contigs, MAGs

The highest and the lowest number of reads belonged to the liquid phase of the sterile feed at 4°C (100 million) and 15°C (64 million), respectively (Supplementary File 1, Table S2). There was statistically significant difference in the number of reads in the biofilm and liquid (Tukey pairwise comparison; P-value = 0.402). We found about 1 million (M) contigs with a total length of nearly 1.5 billion base pair (bp). The largest contig was about 1 Mbp long (Supplementary File 1, Table S3). We recovered about 1519 MAGs. However, only 40 MAGs had at least one putative lipase genes and met the accepted quality threshold to be selected as putative lipolytic MAGs (Supplementary File 1, Table S4).

### 3.2. Lipolytic potential: whole metagenome vs MAGs

Within the metagenomic data, a total of 903 sequences had putative lipolytic activity (EC number 3.1.1.3), of which only 78 were present in putative lipolytic MAGs. By contrast, there were numerous genes coding for the extracellular enzymes that degrade proteins, carbohydrates, short-chain lipids, and phosphates in both the whole metagenome and putative lipolytic MAGs, respectively (The CCR regulatory system for selecting the most suitable carbon source is aligned with the economic theories (Allison and Vitousek 2005). In the presence of simple substrates, cells do not invest carbon and nitrogen for producing extracellular enzymes that decompose complex substrates. However, where carbon and nitrogen resources exist in complex form, producing the relevant enzymes becomes inexpensive (Allison and Vitousek 2005). In the case of lipases, where glucose is abundant, CCR depresses the lipase production (Boekema et al. 2007). In addition, the expression of *proteases affects the lipase production (Andersson 1980, Black and DiRusso 2003). In Bacillus* subtills, for example, the accumulation of amino acids induced the cells to produce more proteases and depress the lipase expression. Table 1).

**Table 1.** Comparison between the number of extracellular hydrolytic enzymes in the whole metagenome and putative lipolytic MAGs.

| Enzyme class | Number in the whole metagenomes | Number in the MAGs |
| --- | --- | --- |
| Proteases | 135,456 | 1,272 |
| Phosphatases | 91,147 | 764 |
| Carbohydrate degraders | 47,893 | 663 |
| Esterases/phospholipases | 6,189 | 200 |
| Lipases | 903 | 78 |

Three most frequently annotated genes for degrading sugars in the whole metagenome were *β-galactosidase, β-glucosidase*, and *α-galactosidase.* However, in the putative lipolytic MAGs, *β-glucosidase, β-hexosaminidase*, *cellulase, α-amylase, α-galactosidase*, and *endo-β-xylanase* were the most annotated. The large difference in the number of the genes can indicate that cells might have various alternative gene regulatory systems for expressing the genes which are involved in degrading sugars rather than lipases. Bacteria have a global regulatory mechanism known as carbon catabolite repression (CCR). In the presence of easily accessible carbon sources like sugars, CCR inhibits the expression of genes that allow cells to use a secondary carbon source (Görke and Stülke 2008). One of the key genes in this process is catabolite repression resistance gene, known as the *phosphotransferase system sugar specific EII component (PTS-EII)* or *putative sugar kinases*. These genes were present in all putative lipolytic MAGs (Supplementary File 1, Table S5).

The CCR regulatory system for selecting the most suitable carbon source is aligned with the economic theories (Allison and Vitousek 2005). In the presence of simple substrates, cells do not invest carbon and nitrogen for producing extracellular enzymes that decompose complex substrates. However, where carbon and nitrogen resources exist in complex form, producing the relevant enzymes becomes inexpensive (Allison and Vitousek 2005). In the case of lipases, where glucose is abundant, CCR depresses the lipase production (Boekema et al. 2007). In addition, the expression of proteases affects the lipase production (Andersson 1980, Black and DiRusso 2003). In *Bacillus subtills*, for example, the accumulation of amino acids induced the cells to produce more proteases and depress the lipase expression.

### 3.3. Which are the lipolytic MAGs and why they have lipases?

Putative lipolytic MAGs belonged to 14 distinct phyla (mostly from the *Actinobacteria*, *Proteobacteria* and *Bacteroidota*), with two unclassified at the phyla level (Figure 1).

**Figure 1.**
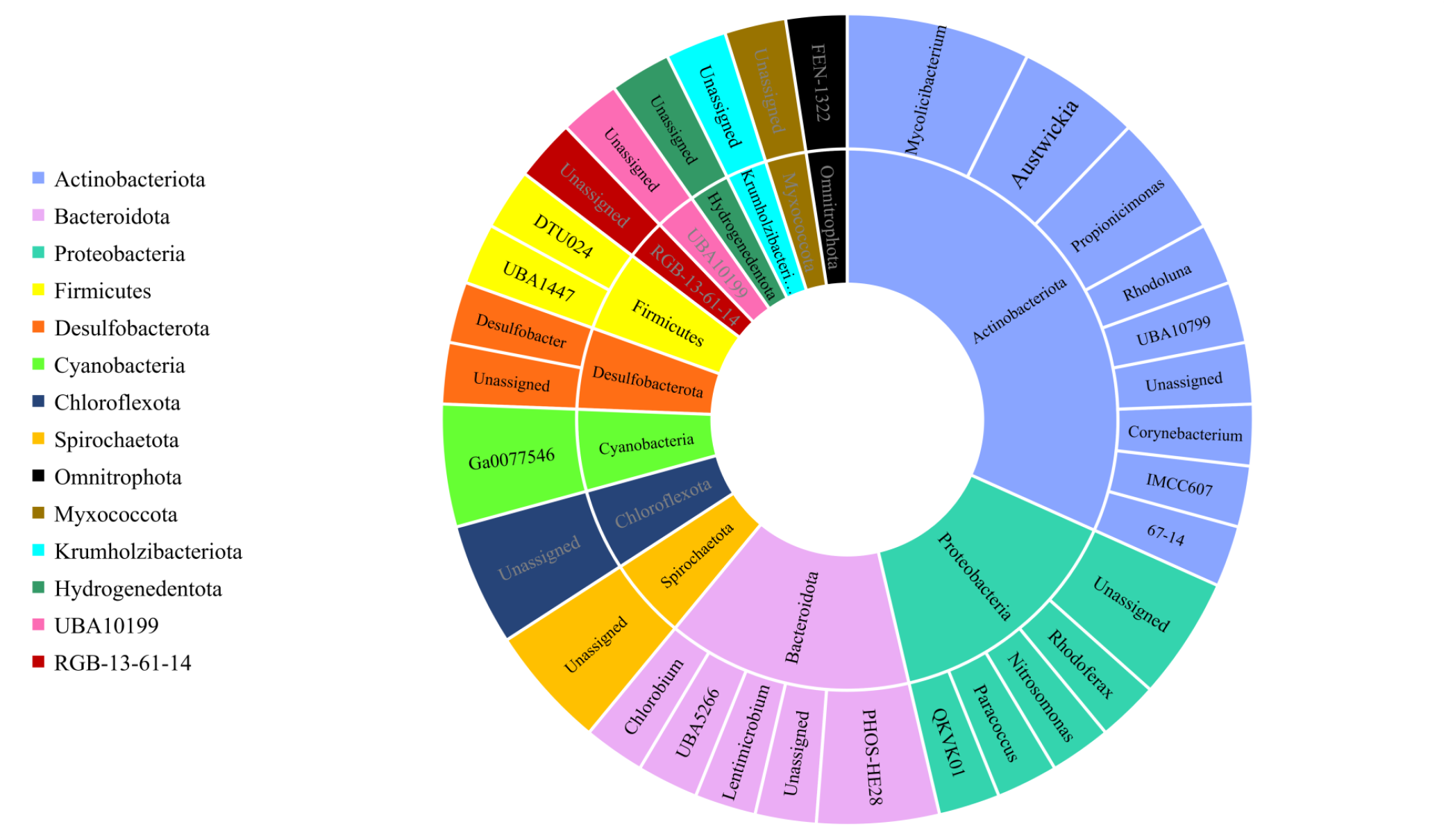
Taxonomic classification of good lipolytic MAGs at phylum and genus level.

Some of the putative lipase genes were found from genera we did not expect to be lipolytic or indeed in anaerobic reactors (such as aerobic autotrophs). We have classified the putative lipolytic MAGs into three categories: i) a possible MAG with a lipase gene but no *fadL* gene to transport long-chain fatty acids; ii) a true lipid degrader: a MAG with both lipase and *fadL* genes; and iii) a miscellaneous lipid degrader: a MAG that degrades lipids for other purposes like denitrification, polyhydroxyalkanoates (PHAs) accumulation/degradation or invasion of other bacteria’s outer membrane (Supplementary File 1, Table S6). The fourth possibility is that these are mis-assemblies or mis-annotations. Even high-quality MAGs can be subjected to these misinterpretations.

We could not label any MAG with certainty as a possible or true lipid degrader due to both non-universality of *fadL* gene and mis-assembly/mis-annotation possibility. For Gram-positive bacteria, still no universal known long-chain fatty acid transporter protein like the *fadL* in Gram-negatives, is characterised (Salvador López and Van Bogaert 2021). Hence, we could not decide which of the putative Gram-positive lipolytic MAGs (13 from the phylum *Actinobacteria* and 2 from the *Firmicutes_A*) are a true lipid degrader. Also, in putative Gram-negative lipolytic MAGs, only 2 out of 18 (*Bin 967* and *Bin 1501*, respectively, represented *Rhodoferax* and an unclassified genus from *Syntrophorhabdia* class in *Desulfobacterota* phylum) had both lipase and *fadL* gene. The absence of *fadL* in the rest of the 16 MAGs might be because of the mis-assembly and mis-annotation.

Additionally, we assumed the co-presence of lipases and other genes in the MAG, like the essential denitrification genes, or genes required for synthesizing or degrading PHAs, might be a sign of miscellaneous lipid degrader.

One of the most curious lipolytic MAGs was *Bin 22*, a possible *Nitrosomonas*. The presence of lipase gene in this genome seemed redundant as *Nitrosomonas* are aerobic nitrifiers, and classically utilize carbon dioxide as a carbon source. However, some species like *Nitrosomonas europaea* are facultative anaerobes and some have even shown denitrification activity under anaerobic conditions. The link between lipolysis and denitrification has been shown in some studies. Denitrifying bacteria utilize long-chain fatty acids in the absence of light in anaerobic reactors, (Mackie et al. 1991) and anaerobic denitrifiers like *Acidovorax caeni sp. nov.* have lipase activity (Heylen et al. 2008).

Besides, PHA production/degradation is linked to lipolysis as well. Bacteria that accumulate PHA, either produce lipases to degrade oily substrates and obtain carbon to store PHA (Tufail et al. 2017) or degrade the intracellular PHA when the carbon is limited (Mitra et al. 2020). *Nitrosomonas* have been proposed as a PHA-producing bacterium (Yang et al. 2013). Previous reports have suggested that many bacteria, including denitrifiers, produce lipases rather than polymerases to degrade PHAs, though the reason is not known (Chu and Wang 2017, Jaeger et al. 1995, Muhammadi et al. 2015, Sharma et al. 2019, Wang and Chu 2016). We hypothesized that either anaerobic condition or the presence of PHA or other bacteria might induce the lipase expression. The assimilation of long-chain fatty acids, such as palmitic acid, in anaerobic conditions represses ammonia-oxidation activity of nitrifiers (Juliette et al. 1995). The presence of global nitrogen regulatory gene (*ntcA*), which existed in *Bin 22*, can activate the assimilation of other nitrogen sources if ammonium/NH_4_^+^ is absent (Lee et al. 1999). Also, when *Nitrosomonas sp. Is79* was co-cultured with *Nitrobacter winogradskyi*, the abundance of periplasmic lipases in its proteome increased (Sedlacek et al. 2016).

One possible explanation for the presence of lipase in the *Bin 22* therefore might be that it represents an uncharacterised facultative *Nitrosomonas* species that use the lipase for denitrification and PHA production/degradation.

Topologically the closest species to *Bin 22* was *Nitrosomonas sp003201565* (Supplementary File 1, Table S7) deposited in the protein database of the *National Center for Biotechnology Information (NCBI*) as *Nitrosomonas sp. Nm84* (accession number: QJJP01000015). This genome, from a pure culture, not only had the lipase and *fadL* genes, but like *Bin 22*, it contained the essential denitrification genes including *nirK* (Copper-containing nitrite reductase), *norB* and *norC* (nitric oxide subunit B and C) (Braker et al. 2000, Torregrosa-Crespo et al. 2017). However, they both lacked the PHA synthesising genes. On the other hand, 13 putative lipolytic MAGs from phyla *Proteobacteria* and *Actinobacteria* had either only PHA synthesizing genes (e.g., *PhaC*) or both PHA synthesizing and denitrification genes (Supplementary File 1, Table S6). Therefore, we assumed that *Bin 22* might use the lipase for degrading the PHA produced by other bacteria from these two phyla for denitrification. Similarly, the other 10 MAGs from several phyla that only had denitrification and lipase genes (no PHA synthesising genes) might use the lipase for degrading the PHA produced by others.

Furthermore, we have searched for potential genes involved in the export of lipases to the extracellular medium in *Bin 22* to validate the presence of lipase genes. Gram-negative bacteria use both *Type I* and *Type II secretion system* for exporting lipases (Ahn et al. 1999, Hausmann and Jaeger 2010). *Type I* secretion pathway usually involves the expression of *ATP-binding Cassette (ABC*) transporters consisted of *ABC* proteins, membrane fusion proteins (MFP) and outer membrane proteins (OMP) at the upstream of the lipase gene. In addition to this, the lipase gene itself should contain several conserved glycine-rich motifs of *GGXGXD* (G, glycine, X, any amino acid, D, Aspartic acid) known as *LAAD*/lipase *ABC* transporter recognition domain at the C-terminal (Chung et al. 2009). Nonetheless, none of the aforementioned exports genes or motifs were found in *Bin 22* or in the associated public genome of *Nitrosomonas sp. Nm84.* Only one of the related lipases (accession number PXW86082) in the public genome had the motifs at the C-terminal.

There were also 16 lipase containing MAGs (those with known genus were all facultative anaerobes) that had no denitrification nor PHA synthesizing genes. For most of them we do not know what the exact role of lipases is. For example, *Chlorobium* in *Bin 803* are photosynthetic green sulphur-reducing bacteria. This MAG, however, had both dark-operative protochlorophyllide reductase (*BChl*) and light-harvesting antenna/chlorosomes (*csmA*) genes that enable *Chlorobium* to survive at extremely low light conditions (Frigaard et al. 2003). Two *Chlorobium* species in *NCBI* had also lipase genes but no *fadL* genes including *Chlorobium limicola* (accession number KUL20464) and *Chlorobium phaeobacteroides DSM 26* (accession number ABL66324). *Desulfobacter postgatei (Bin 481*), a sulphate reducing bacteria in *NCBI* had *fadL* gene but no lipase gene. We do not know whether or not this bacterium is a cheater, but uptake of long-chain fatty acids and improved lipid degradation have been confirmed for other sulphate reducers (Alves et al. 2020, Florentino et al. 2020).

### 3.4. Linking the putative lipolytic MAGs to the reactor conditions and lipases

For most putative lipolytic MAGs, the number of mapped reads per reactor conditions did not vary significantly. However, for a few MAGs, statistically significant differences were observed (Supplementary File 1, Table S8 and Figure S1-S3). For instance, considering only the effect of temperature, at 4°C, only Bin 803 (*Chlorobium*) and at 15°C, Bin 328 (Unclassified *Ga0077546* from Cyanobacteria), Bin 231 (Unassigned from *Chloroflexota*), Bin 154 (Unassigned from *Hydrogenedentota*), and Bin 609 (Unclassified *FEN-1322* from *Omnitrophota*) had noticeably higher number of mapped reads.

About 55% of the lipases were in MAGs from the phylum *Actinobacteriota* of which half distributed within two genera, *Mycolicibacterium* and *Corynebacterium.* Both genera existed at both temperatures, treatment and phase, though the latter was slightly (but not statistically significant) higher in the liquid phase (Supplementary File 1, Table S9).

Regardless of their class/taxonomy lipases from the different MAGs were significantly different in length (pairwise Tukey test, P-value = 0.002). One-way ANOVA (pairwise Tukey test, P-value = 0.467) on the length of individual lipases per phylum showed that *Actinobacteriota* had both the largest (819 aa) and the shortest (180 aa) lipases. In addition, the highest and the lowest average length of the lipases were within the phyla *Actinobacteriota* (399 aa) and *Omnitrophota* (220 aa), respectively (Figure 2).

**Figure 2.**
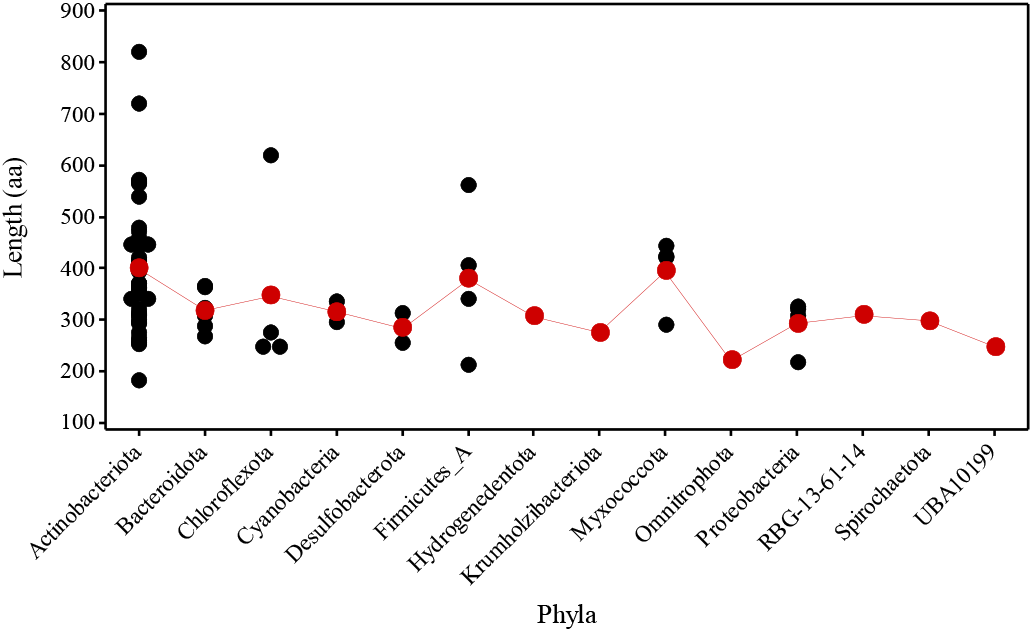
Distribution of the length for individual lipases per phylum 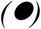 show the length of individual lipases and 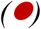 shows the average length of all lipases in a certain phylum (One-way ANOVA, P-value = 0.467).

### 3.5. The putative lipolytic MAGs abundance in the reactors

There were 32 common and abundant (relative abundance ≥ 1%) bacterial genera in at least one reactor conditions (Figure 3).

**Figure 3.**
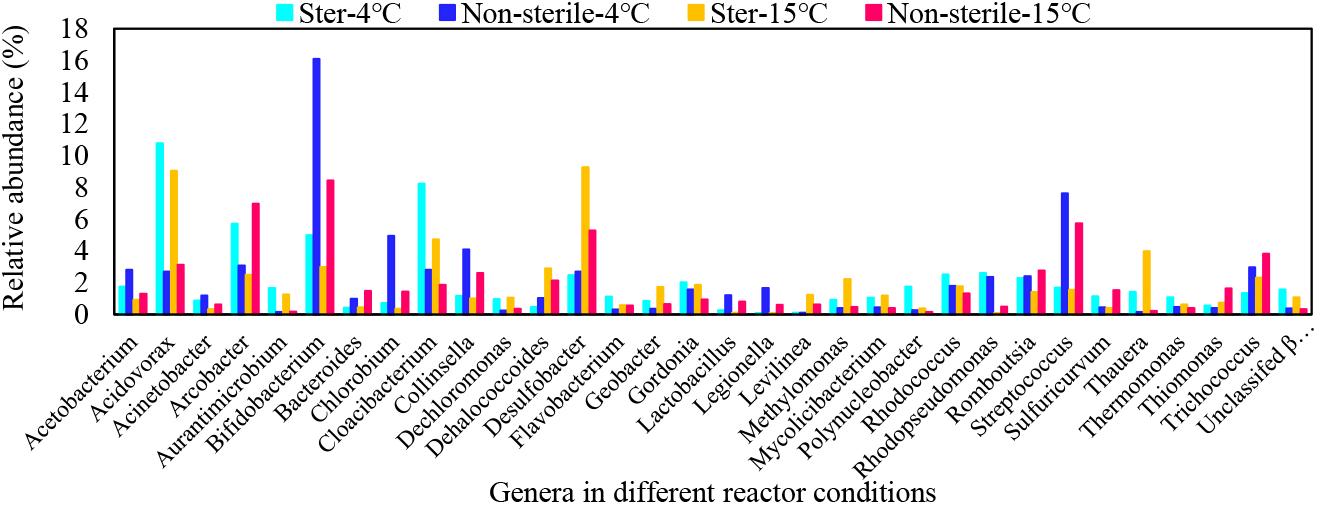
Common genera with more than 1% relative abundance in at least one of the reactor conditions identified by GOTTCHA2.

For most, the effect of temperature and treatment on relative abundance was insignificant (Supplementary File 1, Figure S4). However, *Bifidobacterium* and *Desulfobacter* were more abundant at 4°C and 15°C, respectively. Similarly, a significant effect of treatment was noticeable among the Sterile and Non-sterile fed reactors for *Bifidobacterium*, *Streptococcus*, *Acidovorax* and *Cloacibacterium*. The first two were higher in Non-sterile conditions whereas the second two were the highest at the Sterile conditions.

Only three of the putative lipolytic MAGs (*Desulfobacter*, *Chlorobium* and *Mycolicibacterium*) were both common and abundant (≥1%) in most reactors. *Chlorobium* and *Mycolicibacterium* only had more than 1% abundance at Non-sterile and Sterile conditions, respectively (Figure 3). The rest of the genera identified in MAGs had either very low relative abundance (*Corynebacterium*, *Lentimicrobium*, *Nitrosomonas*, *Paracoccus* and *Rhodoferax*) or they were not represented in the reactors (*Austwickia*, *Propionicimonas* and *Rhodoluna*) (Figure 4).

**Figure 4.**
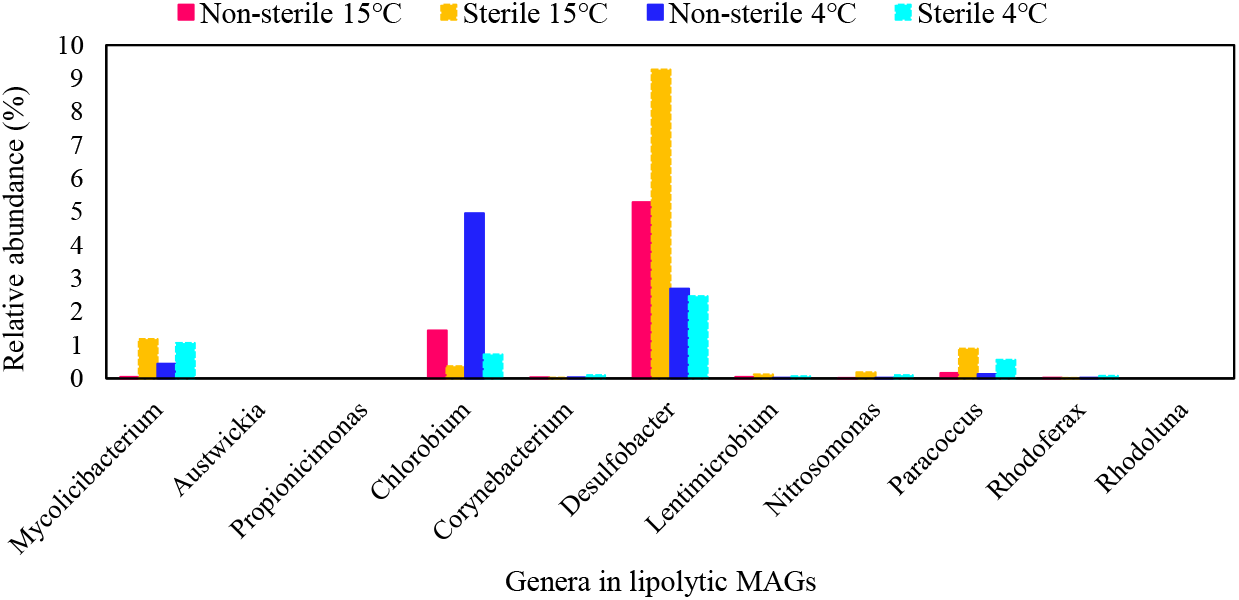
Relative abundance of the genera recovered in MAGs per reactor conditions.

### 3.6. Expressed proteins

A total of 93 and 117 distinct protein groups were found at False Detection Rates (FDR) of 1% and 5%, respectively as listed in Supplementary File 1, Table S10, using the complete metagenomics constructed database. At FDR 5%, there were 24 new protein groups compared to FDR 1% though neither of the new or common hits were significantly different (P-value = 0.514, one-way ANOVA). Not only were none of the hits lipases, none were hydrolytic enzymes of any description either. About 75% of the identified proteins were involved in processing the genetic information, signalling and cellular processes, processing environmental information and energy metabolism. Further 4%, 2%, and 1% of the proteins were related to carbohydrate, amino acids, and lipid metabolism, respectively (Figure 5).

**Figure 5.**
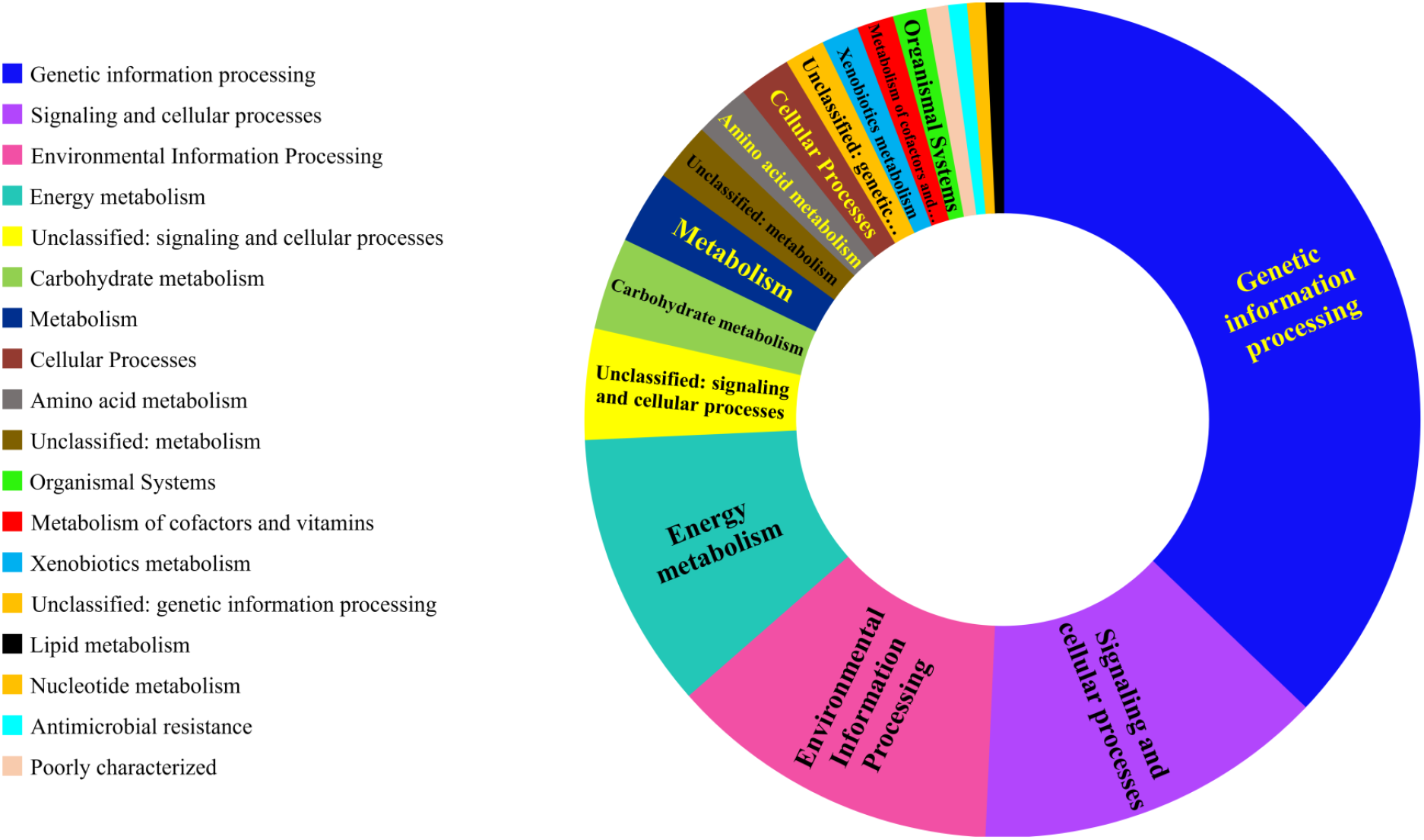
Functional classification of identified proteins at FDR 5% based on KEGG database.

In terms of class, *outer membrane porin proteins (omp32*) outnumbered the rest of the classes (25 %) and after them in descending order there were *vitamin B12 transporters (btuB), TonB-dependent starch-binding receptors (susC*) and *major outer membrane proteins P. IA (porA).*

We detected several *porins*, *ABC* transporters like *lamB* (*Maltoporin*) and *fadL* (long-chain fatty acid transporters), and the latter is of particular interest. The expression of *fadL* might be related to the expression of lipases. We assumed that cells would only invest on expressing *fadL* genes when expressed lipases had already released long-chain fatty acids from the lipidic molecules.

Several cytoplasmic proteins were present including *groEL (60 KDa chaperonin), tufA (elongation factor Tu), fusA (elongation factor G), rpsA (30S ribosomal protein S1), rpsC (30S ribosomal protein S3), rpsE (30S ribosomal protein S5), rpsG (30S ribosomal protein S7*) and *rpsP (30S ribosomal protein S16).* The presence of these proteins in the extracellular polymeric substances (EPS) is related to either the presence of extracellular vesicles in the EPS or cell lysis that happens during the biofilm maturation (Jachlewski et al. 2015).

Therefore, we profiled several proteins that are typically found in the extracellular vesicles. These proteins were outer membrane proteins and *porins* (*ompA, ompW, ompX ompF, porA* and *porB*), proteins that release toxic compounds and attack the competing bacteria (*acrA* or *multidrug efflux pump subunit*), nutrient sensors and transporters that carry certain molecules under nutrient limited conditions like *ABC transporters (fadL, lamB, btuB*) and *TonB-dependent receptors (susC*). Particularly, the presence of *fadL* genes in the extracellular vesicles is interesting. They were frequently found as the top 50 vesicular proteins in Gram-negative bacteria (Lee et al. 2016). Nonetheless, their exact role and why they are there is yet to be known.

### 3.7. Taxonomical distribution of identified proteins by metaproteomics

About 97% of the identified expressed genes (at FDR 5%) were related to the bacterial domain and at least from 19 distinct classes (Figure 6a), among which *Betaproteobacteria* had the greatest share (57 %). The top-ranked identified genera with expressed proteins were all from the class *Betaproteobacteria* including *Paucimonas*, *Dechloromonas*, *Acidovorax*, *Azoarcus* and *Thauera*, respectively (Figure 6b). The full list of all genera associated to the expressed proteins is presented in Supplementary File 1 Table S11. Among the top-ranked, all genera except for *Paucimonas* have been formerly identified by *GOTTCHA2* and their relative abundance in each reactor was known (Figure 7). Comparatively, *Azoarcus*, was the only low-abundant genus with no abundance at Non-sterile-4°C.

**Figure 6.**
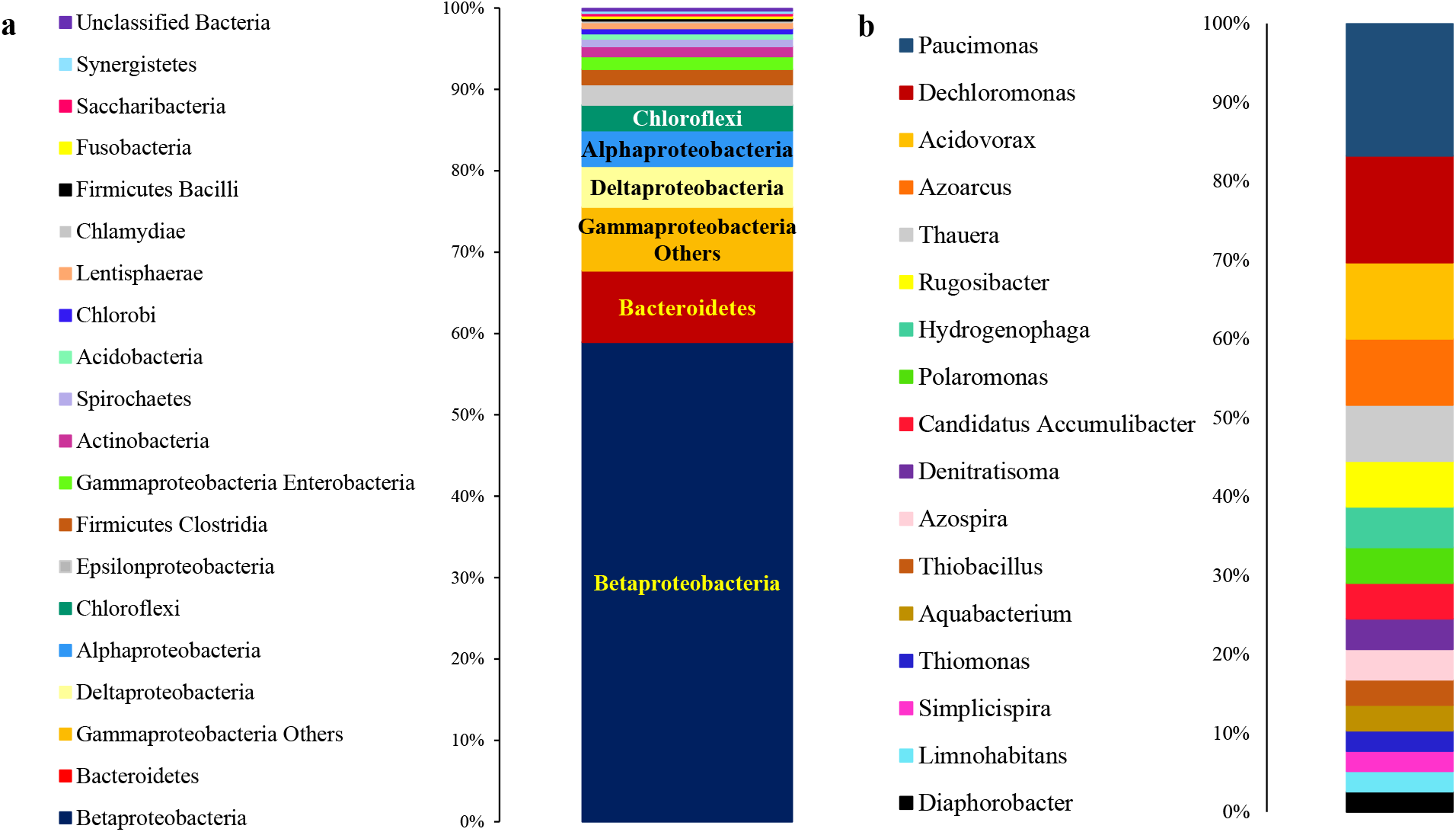
a) Taxonomic distribution of expressed proteins at class level; b) list of genera that had more than three expressed proteins (FDR 5 %).

**Figure 7.**
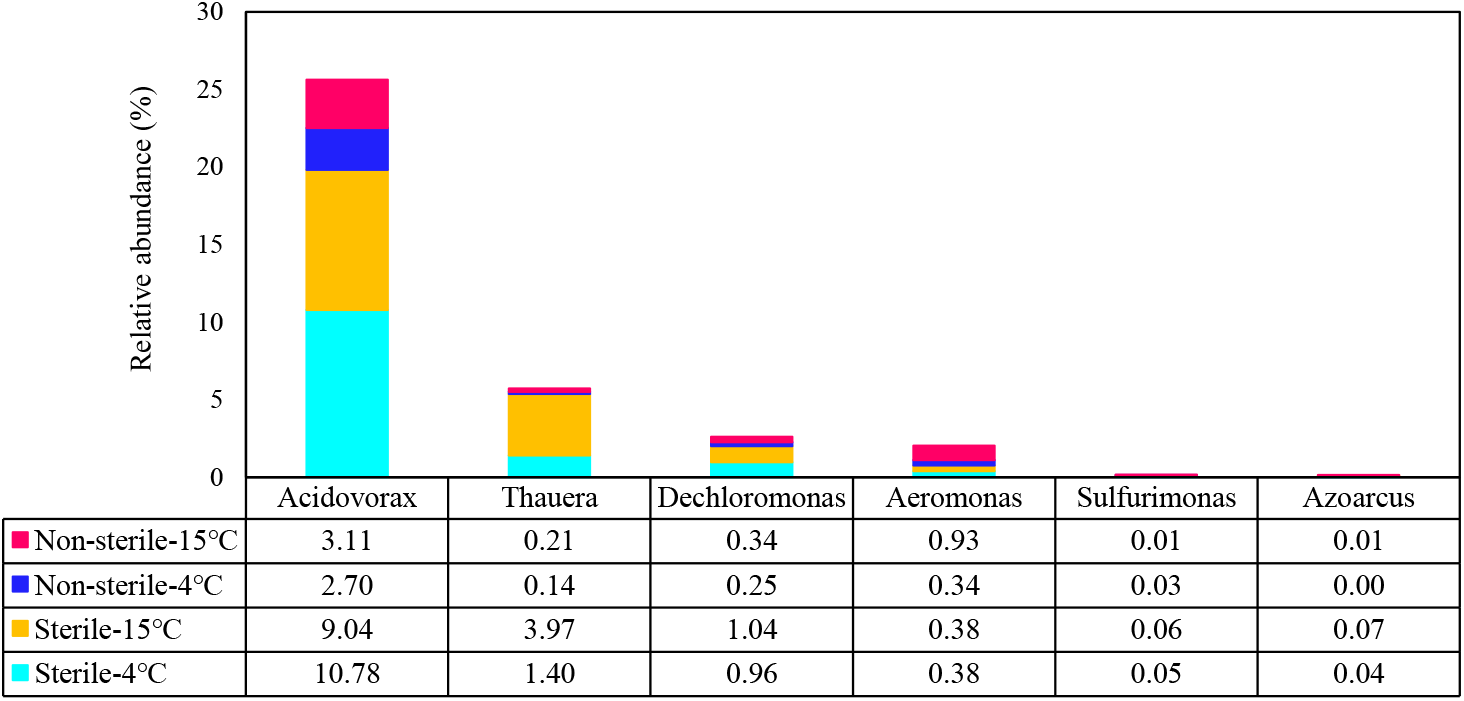
Relative abundance of top-ranked genera per reactors

Although *Paucimonas* was absent from the reactors based on *GOTTCHA2*, it had the highest number of protein matches (Supplementary File 1, Table S12) with most of them being ribosomal proteins or involved in energy metabolism. One *porin* and one *outer membrane protein* were present too.

By contrast, 68% of the related proteins to other genera were *porins* and *outer membrane proteins* (Supplementary File 1, Figure S5). Additionally, both *Dechloromonas* and *Azoarcus* had *fadL* and thus were potentially lipolytic. Expressed *fadL* was also found in two other genera (not among the top-ranked), *Aeromonas* and *Sulfurimonas*. Their relative abundance is presented in Figure 7. *Azoarcus* and *Sulfurimonas* were low-abundant, in all conditions, whereas *Aeromonas* had higher relative abundance (~ 1%) at Non-sterile-15°C.

The expression of *fadL* in *Dechloromonas*, *Azoarcus*, *Aeromonas* and *Sulfurimonas* might imply the presence of long-chain fatty acids in the system and therefore can be a proxy for lipolysis performed by these genera or others. However, none of these four genera were recovered as putative lipolytic MAGs by metagenomics. The absence of lipases along with the presence of *fadL* genes in a genome might be indicative of cheating mechanisms. Nonetheless, the complete genome of these four genera in *NCBI* had both the *fadL* and lipase genes. While this might remove the “cheating label”, from these genera, it does not necessarily make them true lipase producers either. We do not know whether or not *fadL* and lipases are coregulated, but we do know that both can be exported through extracellular vesicles in Gram-negative and Gram-positive bacteria (Galka et al. 2008, Lee et al. 2009, Lee et al. 2016). The presence of both *fadL* and lipases in the extracellular vesicles might have an entirely different reason than the lipolysis of exogenous lipid molecules. For instance, Galka et al. (2008) have shown that pathogens transport lipases as a virulence factor through extracellular vesicles to attack the lipidic membrane of the host cell and deliver lipids to them. The same scenario might apply to bacterial cells interaction, but no study has shown this yet.

### 3.8. Identified proteins of abundant genera

Out of the 32 common bacterial genera with relative abundance of more than 1% (Figure 3) metaproteomics identified proteins expressed by 15 of them (Table 2). More than half (55%) of the proteins were outer membrane proteins and porins. Some of these genera accumulate lipids, e.g., polyhydroxyalkanoates (PHAs) or perform denitrification. Lipid-accumulation is a barrier for lipid degradation in wastewater systems (Chipasa and Mdrzycka 2008) and cold temperature is a stimulator for PHA accumulation (Srivastava et al. 2020).

**Table 2.**
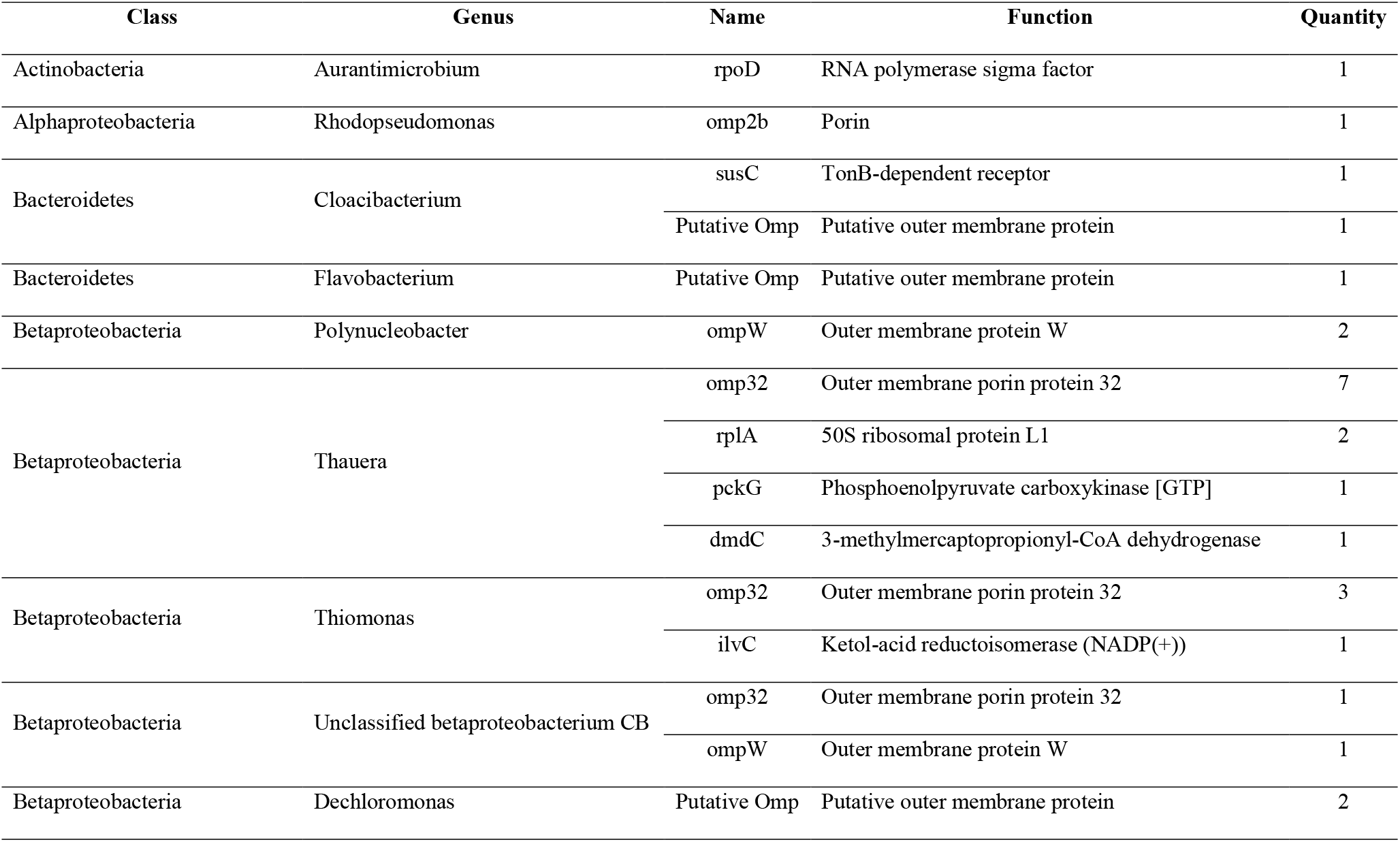
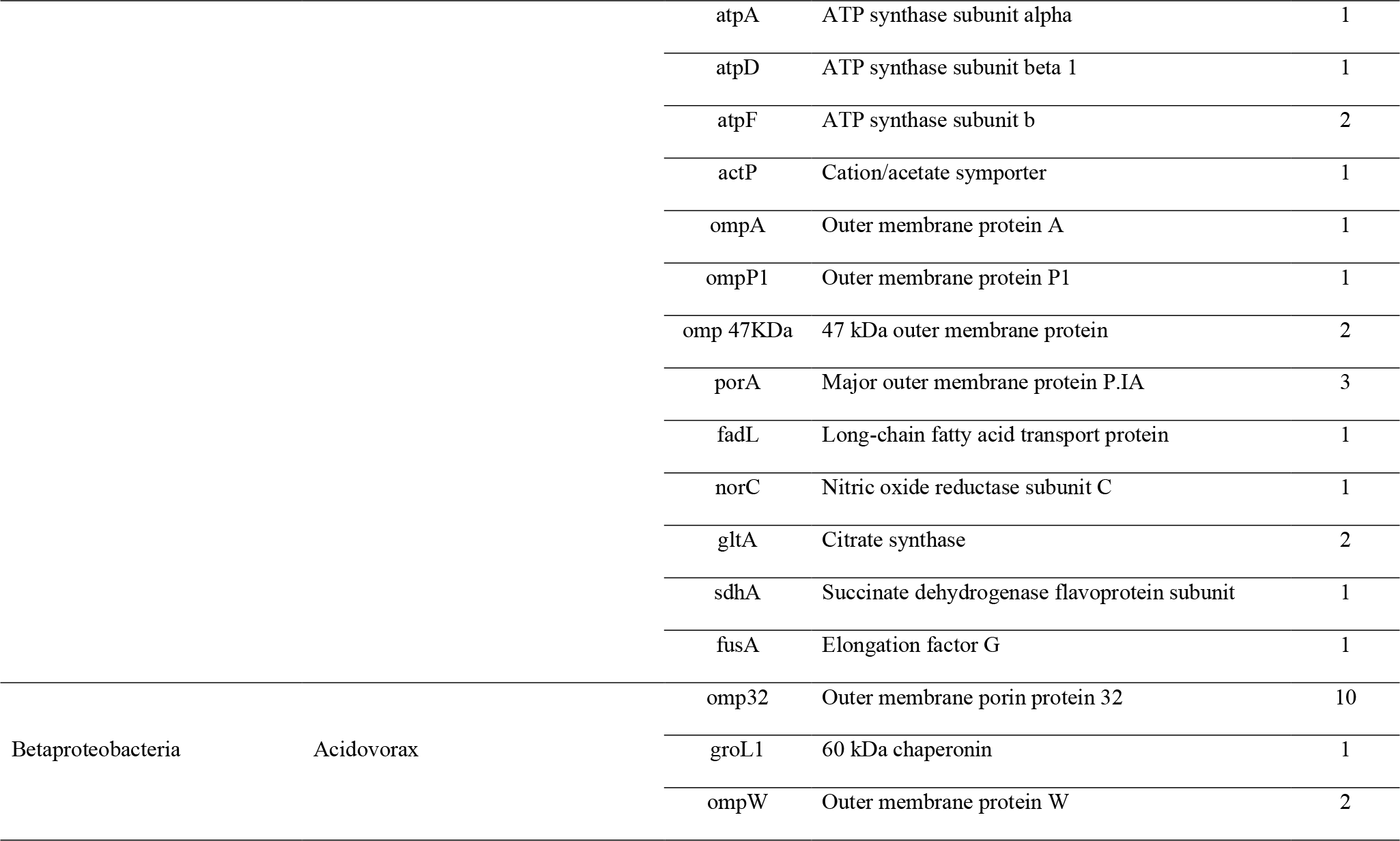
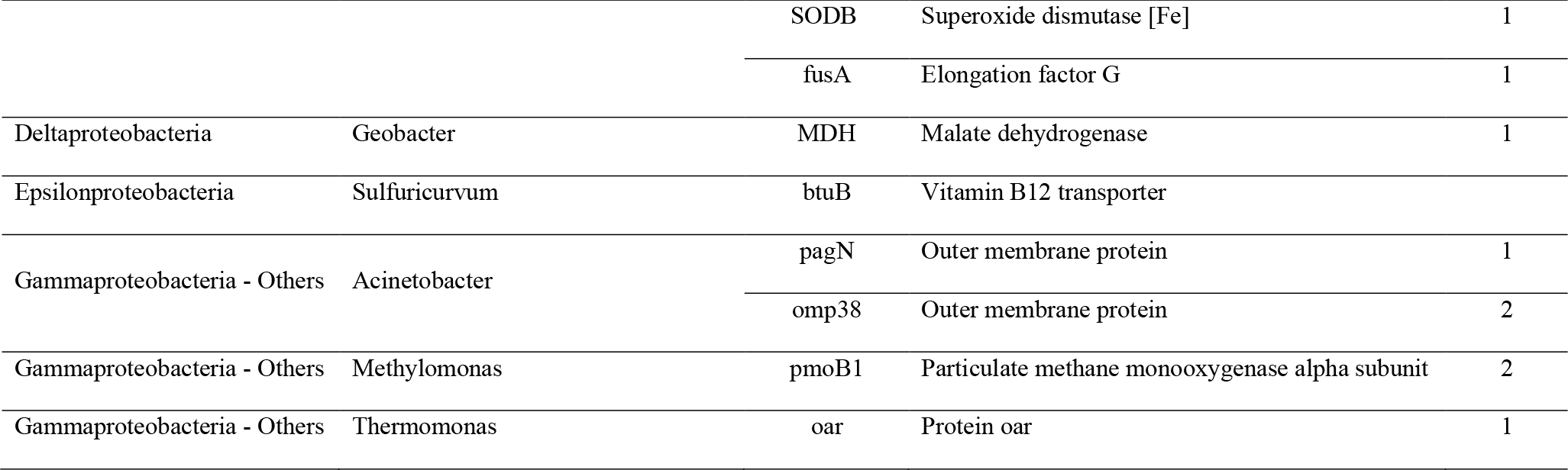
Expressed proteins found from the common genera (?1% relative abundance) per reactor conditions in Figure 3.

At least six of the identified genera (*Acinetobacter*, *Cloacibacterium*, *Dechloromonas*, *Rhodopseudomonas*, *Thauera*, and *Thermomonas*) are involved in PHA accumulation (Carlozzi and Sacchi 2001, Coats et al. 2016, Hauschild et al. 2017, Oshiki et al. 2008, Ram et al. 2018, Singleton et al. 2021). However, *Dechloromonas* was the only genus that had expressed *fadL* gene (but had no lipases). This genus has been identified as an anaerobic denitrifier too (Singleton et al. 2021). The identified *norC* and *actP* (Cation/acetate symporter) proteins for this genus confirm its denitrification activity and competition with methanogens to assimilate acetate (Table 2). Since the complete genome of *Dechloromonas denitrificans* in *NCBI* had lipase, *fadL*, and *phaC* genes (accession numbers respectively, KXB32756 and KXB32487), we assumed that the identified *fadL* of *Dechloromonas* might indicate of its lipolytic activity related to PHA accumulation and denitrification.

Also, the presence of sulphur-reducing bacteria, *Sulfuricurvum* (Table 2) along with the sulphate-reducers, e.g. *Desulfobacter*, has been associated with the occurrence of internal sulphur cycle in the system (St. James and Richardson 2020). Although *Desulfobacter* was not identified by metaproteomics, it was recovered as a good lipolytic MAG and had high abundance at all reactor conditions (Figure 3). Sulphate reduction limits PHA-accumulation, and sulphate-reducers in the absence of sulphate can switch to syntrophic and fermentative metabolisms (St. James and Richardson 2020).

Overall, the combination of metagenomics and metaproteomics failed to identify true lipolytic psychrophiles. Although biomarkers like *fadL* gene were found by metaproteomics, the absence of lipases in the analysed metaproteome prevented any clear conclusion. None of the recovered putative lipolytic MAGs were among the taxa found by metaproteomics. Therefore, we could not determine which cold-adapted microorganism is a true lipid degrader and which might use the lipase for miscellaneous purposes like the PHA accumulation/degradation, denitrification or invasion.

Future studies can therefore investigate how lipolysis and PHA accumulation/degradation are linked and why and when microbes might use lipases rather than PHA polymerases. For more conclusive results, we suggest that some barriers need to be removed by different research disciplines. We can reduce the effect of false positive results in the assembly and annotation steps of metagenomics by integrating both short and long sequenced reads. In metaproteomics on the other hand, we need to develop universal unbiased extraction methods particularly for extracellular enzymes which are more sensitive. Characterising the extracellular vesicle proteins and differentiating them from the EPS proteins is another important issue. Some extracellular proteins like lipases can be present in both location but cells regulate them for different purposes. This can be checked by using synthetic biology techniques that enable determination of the gene regulation mechanisms and the extracellular excretion pathways. Employing high-resolution mass spectrometers along with developing better tools for identifying the mass spectra and matching them to protein groups can also enhance metaproteomics data analysis. And finally, the dependency of metaproteomics to metagenomics database can be reduced through *de novo* approaches.

## 4. Conclusion

- 40 putative lipolytic MAGs were recovered from the metagenomics data of all reactors.
- Extremely lower number of lipases (compared to other hydrolytic extracellular enzymes) were found within the metagenomes.
- Lipolysis may not always occur exogenously. Lipolysis molecules may be linked to PHA accumulation/degradation, denitrification and invasion.
- Most lipases were from phyla *Actinobacteria* and genera *Mycolicibacterium* and *Corynebacterium* that accumulate PHAs.
- Of the 32 common abundant genera in all reactors (relative abundance ≥1%) only three (*Chlorobium*, *Desulfobacter*, and *Mycolicibacterium*) were recovered as putative lipolytic MAGs.
- With few exceptions, there was no significant correlation between the reactor conditions and the number of reads mapped to the putative lipolytic MAGs.
- Out of the 32 common genera profiled by metagenomics, 15 were identified by metaproteomics too; at least 6 of them were involved in lipid/PHA accumulation.
- Metaproteomics identified *fadL* genes for four genera (*Dechloromonas, Azoarcus, Aeromonas* and *Sulfurimonas*), but did not identify any lipases for them. Also, none of these four genera were recovered as putative lipolytic MAGs by metagenomics.
- The proteins identified by metaproteomics were mainly porins and outer membrane proteins and some cytoplasmic proteins were identified too that might enter the EPS through extracellular vesicles.

## Acknowledgements

TPC and IDO acknowledge the support of the EPSRC Frontier Grant A New Frontier in Design: The Simulation of Open Engineered Biological Systems, led by Newcastle University, ref EP/K039083/1.

## Nomenclature

AnMBRs: Anaerobic membrane bioreactors
Bp: Base pair
CCR: Carbon catabolite repression
COD: Chemical oxygen demand
EPS: Extracellular polymeric substances
fadL: Long-chain fatty acid transporter
FDR: False discovery rates
MAGs: Metagenome-assembled genomes
PHAs: Polyhydroxyalkanoates
VSS: Volatile suspended solid

## Notes

### Competing Interest Statement

The authors have declared no competing interest.

