## Supplementary-File 1 for "Looking for lipases and lipolytic organisms in low-temperature anaerobic reactors treating domestic wastewater"

#### Table S1

*Table 1. List of hydrolytic enzymes that was searched against metagenomics data and good lipolytic MAGs*

| Class | Name | EC number | Class | Name | EC number |
|---|---|---|---|---|---|
| Carbohydrate degrading enzymes | β-galactosidase | 3.2.1.23 | Lipolytic enzymes | Triacylglycerol lipase | 3.1.1.3 |
| | β-glucosidase | 3.2.1.21 | | Carboxylesterase | 3.1.1.1 |
| | α-galactosidase | 3.2.1.22 | | Acylglycerol lipase | 3.1.1.23 |
| | β-hexosaminidase | 3.2.1.52 | | Putative phospholipase | 3.1.1.32 |
| | non-reducing end α-L-arabinofuranosidase | 3.2.1.55 | | Lipoprotein lipase | 3.1.1.34 |
| | lysozyme | 3.2.1.17 | | Phospholipase A2 | 3.1.1.4 |
| | 6-phospho-β-glucosidase | 3.2.1.86 | | Lysophospholipase | 3.1.1.5 |
| | Endo-β-xylanase | 3.2.1.8 | | Phosphatidate phoshphohydrolase | 3.1.3.4 |
| | α-amylase | 3.2.1.1 | | Phospholipase C | 3.1.4.3 |
| | Cellulase | 3.2.1.4 | | Phospholipase D | 3.1.4.4 |
| | oligo-1,6-glucosidase | 3.2.1.10 | Proteases | leucyl aminopeptidase | 3.4.11.1 |
| | non-reducing end β-L-arabinofuranosidase | 3.2.1.185 | | enteropeptidase | 3.4.21.9 |
| | α,α-trehalase | 3.2.1.28 | | endopeptidase Clp | 3.4.21.92 |
| | α-D-xyloside xylohydrolase | 3.2.1.177 | | thrombin | 3.4.21.5 |
| | licheninase | 3.2.1.73 | | endopeptidase La | 3.4.21.53 |
| | xylan 1,4-β-xylosidase | 3.2.1.37 | | repressor LexA | 3.4.21.88 |
| | exo-α-sialidase | 3.2.1.18 | | methionyl aminopeptidase | 3.4.11.18 |
| | α-N-acetylgalactosaminidase | 3.2.1.49 | | chymotrypsin | 3.4.21.1 |
| | Levanase | 3.2.1.80 | | serine-type D-Ala-D-Ala carboxypeptidase | 3.4.16.4 |
| | glucan 1,4-β-glucosidase | 3.2.1.74 | | acrosin | 3.4.21.10 |
| | neopullulanase | 3.2.1.135 | | gastricsin | 3.4.23.3 |
| | α-glucosidase | 3.2.1.20 | | signal peptidase II | 3.4.23.36 |

| Class | Name | EC number | Class | Name | EC number |
|---|---|---|---|---|---|
| | UDP-N,N'-diacetylbacillosamine 2-epimerase (hydrolysing) | 3.2.1.184 | | signal peptidase I | 3.4.21.89 |
| | unsaturated rhamnogalacturonyl hydrolase | 3.2.1.172 | | peptidase Do | 3.4.21.107 |
| **Carbohydrate degrading enzymes** | 6-phospho-β-galactosidase | 3.2.1.85 | **Proteases** | tripeptide aminopeptidase | 3.4.11.4 |
| | exo-1,4-β-D-glucosaminidase | 3.2.1.165 | | carboxypeptidase A | 3.4.17.1 |
| | galacturan 1,4-α-galacturonidase | 3.2.1.67 | | Xaa-Pro aminopeptidase | 3.4.11.9 |
| | Mannosylglycerate hydrolase | 3.2.1.170 | | membrane alanyl aminopeptidase | 3.4.11.2 |
| | β-glucuronidase | 3.2.1.31 | | D-stereospecific aminopeptidase | 3.4.11.19 |
| | glucan 1,4-α-glucosidase | 3.2.1.3 | | interstitial collagenase | 3.4.24.7 |
| | β-fructofuranosidase | 3.2.1.26 | | HslU—HslV peptidase | 3.4.25.2 |
| | cyclomaltodextrinase | 3.2.1.54 | | cytosol nonspecific dipeptidase | 3.4.13.18 |
| | pullulanase | 3.2.1.41 | | oligopeptidase A | 3.4.24.70 |
| | mannan endo-1,4-β-mannosidase | 3.2.1.78 | | Xaa-Pro dipeptidase | 3.4.13.9 |
| | Endo-β-glucosidase | 3.2.1.39 | | C-terminal processing peptidase | 3.4.21.102 |
| | chitinase | 3.2.1.14 | | carboxypeptidase Taq | 3.4.17.19 |
| | 4-α-D-{(1→4)-α-D-glucano}trehalose trehalohydrolase | 3.2.1.141 | | D-Ala-D-Ala dipeptidase | 3.4.13.22 |
| | sulfoquinovosidase | 3.2.1.199 | | prolyl aminopeptidase | 3.4.11.5 |
| | cellulose 1,4-β-cellobiosidase (non-reducing end) | 3.2.1.91 | | acylaminoacyl-peptidase | 3.4.19.1 |
| | gellan tetrasaccharide unsaturated glucuronyl hydrolase | 3.2.1.179 | | chymotrypsin C | 3.4.21.2 |
| | unsaturated chondroitin disaccharide hydrolase | 3.2.1.180 | | prolyl oligopeptidase | 3.4.21.26 |
| | xylan α-1,2-glucuronosidase | 3.2.1.131 | | rhomboid protease | 3.4.21.105 |
| | Arabinosidase | 3.2.1.99 | | dipeptidyl-peptidase I | 3.4.14.1 |
| | xyloglucan-specific endo-beta-1,4-glucanase | 3.2.1.151 | | glutathione γ-glutamate hydrolase | 3.4.19.13 |
| | keratan-sulfate endo-1,4-β-galactosidase | 3.2.1.103 | | peptidyl-dipeptidase Dcp | 3.4.15.5 |
| | UDP-N-acetylglucosamine 2-epimerase (hydrolysing) | 3.2.1.183 | | dipeptidyl-peptidase IV | 3.4.14.5 |
| | α,α-phosphotrehalase | 3.2.1.93 | | bacterial leucyl aminopeptidase | 3.4.11.10 |
| | arabinogalactan endo-β-1,4-galactanase | 3.2.1.89 | | dipeptidase E | 3.4.13.21 |
| | oligosaccharide reducing-end xylanase | 3.2.1.156 | | β-peptidyl aminopeptidase | 3.4.11.25 |
| | glucan 1,3-β-glucosidase | 3.2.1.58 | | prolyltripeptidyl aminopeptidase | 3.4.14.12 |

| Class | Name | EC number | Class | Name | EC number |
|---|---|---|---|---|---|
| | β-porphyranase | 3.2.1.178 | **Proteases** | β-aspartyl-peptidase | 3.4.19.5 |
| | glucan 1,4-α-maltohexaosidase | 3.2.1.98 | | oligopeptidase B | 3.4.21.83 |
| **Carbohydrate degrading enzymes** | maltose-6'-phosphate glucosidase | 3.2.1.122 | | γ-D-glutamyl-L-lysine dipeptidyl-peptidase | 3.4.14.13 |
| | chitosanase | 3.2.1.132 | | glutamate carboxypeptidase | 3.4.17.11 |
| | xylan 1,3-β-xylosidase | 3.2.1.72 | | pyroglutamyl-peptidase I | 3.4.19.3 |
| | (Ara-f)3-Hyp β-L-arabinobiosidase | 3.2.1.187 | | proteasome endopeptidase complex | 3.4.25.1 |
| | κ-carrageenase | 3.2.1.83 | | coagulation factor Xa | 3.4.21.6 |
| | limit dextrin α-1,6-maltotetraose-hydrolase | 3.2.1.196 | | bleomycin hydrolase | 3.4.22.40 |
| | endo-polygalacturonase | 3.2.1.15 | | arginyl aminopeptidase | 3.4.11.6 |
| | glucan 1,6-α-glucosidase | 3.2.1.69 | | cathepsin D | 3.4.23.5 |
| | exo-poly-α-galacturonosidase | 3.2.1.82 | | HycI peptidase | 3.4.23.51 |
| | glucan 1,4-α-maltotetraohydrolase | 3.2.1.60 | | subtilisin | 3.4.21.62 |
| | isoamylase | 3.2.1.68 | | cyanophycinase | 3.4.15.6 |
| | glucuronoarabinoxylan endo-1,4-β-xylanase | 3.2.1.136 | | muramoyltetrapeptide carboxypeptidase | 3.4.17.13 |
| | protein O-GlcNAcase | 3.2.1.169 | | lysostaphin | 3.4.24.75 |
| | λ-carrageenase | 3.2.1.162 | | gpr endopeptidase | 3.4.24.78 |
| | α-agarase | 3.2.1.158 | | γ-D-glutamyl-meso-diaminopimelate peptidase I | 3.4.19.11 |
| | mannosyl-glycoprotein endo-β-N-acetylglucosaminidase | 3.2.1.96 | | SpoIVB peptidase | 3.4.21.116 |
| | endo-1,3-β-xylanase | 3.2.1.32 | | Xaa-Pro dipeptidyl-peptidase | 3.4.14.11 |
| | glucan 1,6-α-isomaltosidase | 3.2.1.94 | | ficain | 3.4.22.3 |
| | glucan 1,4-α-maltohydrolase | 3.2.1.133 | | gingipain R | 3.4.22.37 |
| | 2,6-β-fructan 6-levanbiohydrolase | 3.2.1.64 | | carboxypeptidase T | 3.4.17.18 |
| | dextranase | 3.2.1.11 | | lysyl endopeptidase | 3.4.21.50 |
| | β-agarase | 3.2.1.81 | | serralysin | 3.4.24.40 |
| | endo-α-N-acetylgalactosaminidase | 3.2.1.97 | | glutamyl aminopeptidase | 3.4.11.7 |
| | hyaluronoglucosaminidase | 3.2.1.35 | | clostripain | 3.4.22.8 |
| | blood-group-substance endo-1,4-β-galactosidase | 3.2.1.102 | | pitrilysin | 3.4.24.55 |

| Class | Name | EC number | Class | Name | EC number |
|---|---|---|---|---|---|
| | ι-carrageenase | 3.2.1.157 | | chymosin | 3.4.23.4 |
| | β-amylase | 3.2.1.2 | | aminopeptidase S | 3.4.11.24 |
| **Phosphatase** | Alkaline phosphatase | 3.1.3.1 | **Proteases** | plasminogen activator Pla | 3.4.23.48 |
| | Acid phosphatase | 3.1.3.2 | | gingipain K | 3.4.22.47 |
| | Phosphoserine phosphatase | 3.1.3.3 | | zinc D-Ala-D-Ala carboxypeptidase | 3.4.17.14 |
| | Phosphatidate phosphatase | 3.1.3.4 | | trypsin | 3.4.21.4 |
| | 5'-nucleotidase | 3.1.3.5 | | PepB aminopeptidase | 3.4.11.23 |
| | 3'-nucleotidase | 3.1.3.6 | | sedolisin | 3.4.21.100 |
| | 3'(2'),5'-bisphosphate nucleotidase | 3.1.3.7 | | glutamyl endopeptidase | 3.4.21.19 |
| | 3-phytase | 3.1.3.8 | | thermitase | 3.4.21.66 |
| | Glucose-6-phosphatase | 3.1.3.9 | | bacillolysin | 3.4.24.28 |
| | Glucose-1-phosphatase | 3.1.3.10 | | xanthomonalisin | 3.4.21.101 |
| | Fructose-bisphosphatase | 3.1.3.11 | | cathepsin B | 3.4.22.1 |
| | Trehalose-phosphatase | 3.1.3.12 | | streptopain | 3.4.22.10 |
| | Histidinol-phosphatase | 3.1.3.15 | | thermolysin | 3.4.24.27 |
| | Protein-serine/threonine phosphatase | 3.1.3.16 | | aqualysin 1 | 3.4.21.111 |
| | Phosphoglycolate phosphatase | 3.1.3.18 | | microbial collagenase | 3.4.24.3 |
| | Glycerol-1-phosphatase | 3.1.3.21 | | pseudolysin | 3.4.24.26 |
| | Mannitol-1-phosphatase | 3.1.3.22 | | α-Lytic endopeptidase | 3.4.21.12 |
| | Sugar-phosphatase | 3.1.3.23 | | lactocepin | 3.4.21.96 |
| | Inositol-phosphate phosphatase | 3.1.3.25 | | flavastacin | 3.4.24.76 |
| | Phosphatidylglycerophosphatase | 3.1.3.27 | | atrolysin A | 3.4.24.1 |
| | 3-deoxy-manno-octulosonate-8-phosphatase | 3.1.3.45 | | IgA-specific metalloendopeptidase | 3.4.24.13 |
| | Protein-tyrosine-phosphatase | 3.1.3.48 | | C5a peptidase | 3.4.21.110 |
| | Phosphatidylinositol-3-phosphatase | 3.1.3.64 | | omptin | 3.4.23.49 |
| | 2-deoxyglucose-6-phosphatase | 3.1.3.68 | | vibriolysin | 3.4.24.25 |
| | Mannosyl-3-phosphoglycerate phosphatase | 3.1.3.70 | | Pro-Pro endopeptidase | 3.4.24.89 |
| | 2-phosphosulfolactate phosphatase | 3.1.3.71 | | aureolysin | 3.4.24.29 |

| Class | Name | EC number | Class | Name | EC number |
|---|---|---|---|---|---|
| | Adenosylcobalamin/alpha-ribazole phosphatase | 3.1.3.73 | | streptogrisin B | 3.4.21.81 |
| | Pyridoxal phosphatase | 3.1.3.74 | | β-lytic metalloendopeptidase | 3.4.24.32 |
| Phosphatase | Acireductone synthase | 3.1.3.77 | Proteases | snapalysin | 3.4.24.77 |
| | Phosphatidylinositol-4,5-bisphosphate 4-phosphatase | 3.1.3.78 | | streptogrisin A | 3.4.21.80 |
| | Mannosylfructose-phosphate phosphatase | 3.1.3.79 | | mycolysin | 3.4.24.31 |
| | D-glycero-beta-D-manno-heptose 1,7-bisphosphate 7-phosphatase | 3.1.3.82 | | | |
| | D-glycero-alpha-D-manno-heptose-1,7-bisphosphate 7-phosphatase | 3.1.3.83 | | | |
| | Glucosyl-3-phosphoglycerate phosphatase | 3.1.3.85 | | | |
| | 87 2-hydroxy-3-keto-5-methylthiopentenyl-1-phosphate phosphatase | 3.1.3.87 | | | |
| | 89 5'-deoxynucleotidase | 3.1.3.89 | | | |
| | 90 Maltose 6'-phosphate phosphatase | 3.1.3.90 | | | |
| | 97 3',5'-nucleoside bisphosphate phosphatase | 3.1.3.97 | | | |
| | 101 Validoxylamine A 7'-phosphate phosphatase | 3.1.3.101 | | | |
| | 104 5-amino-6-(5-phospho-D-ribitylamino)uracil phosphatase | 3.1.3.104 | | | |

### Table S2

*Table 2. Reads generated after sequencing the DNA extractes of both liquid and biofilm phase in AnMBRs.*

| Sample | Number of generated reads | |
|---|---|---|
| | Liquid phase | Biofilm phase |
| 4℃-Nster 1 | 78,901,230 | 85,281,044 |
| 4℃-Nster 2 | 78,902,998 | 82,749,706 |
| 4℃-Ster 1 | 100,225,412 | 83,599,124 |
| 4℃-Ster 2 | 86,566,622 | 65,788,040 |
| 15℃-Nster 1 | 73,358,304 | 92,205,090 |
| 15℃-Nster 2 | 71,669,572 | 81,414,448 |
| 15℃-Ster 1 | 76,591,032 | 82,766,216 |
| 15℃-Ster 2 | 64,788,634 | 89,683,802 |

### Table S3

*Table 3. Contigs statistics obtained from the co-assembly of the reads of the AnMBRs.*

| Contigs information | Statistics |
|---|---|
| Total number of contigs | 1,109,690 |
| # contigs (>= 0 bp) | 1,109,690 |
| # contigs (>= 1,000 bp) | 352,375 |
| # contigs (>= 10,000 bp) | 9,103 |
| # contigs (>= 100,000 bp) | 142 |
| # contigs (>= 1,000,000 bp) | 1 |
| Largest contig (bp) | 1,226,853 |
| Total length (bp) | 1,428,194,318 |
| Total length (>= 0 bp) | 1,428,194,318 |
| Total length (>= 1000 bp) | 913,963,731 |
| Total length (>= 10000 bp) | 210,083,869 |
| Total length (>= 100000 bp) | 25,407,583 |
| Total length (>= 1000000 bp) | 1,226,853 |
| N50 (bp) | 1,490 |
| N75 (bp) | 782 |
| L50 | 186,044 |
| L75 | 531,442 |
| GC (%) | 52.49 |

### Table S4

*Table 4. Bacteria marker summary used by GTDB per MAGs and the contamination and genome completeness information of the MAGs.*

| Name | Unique Gene Count | Multiple Gene Count | Missing Gene Count | Genome completeness (%) | Contamination (%) |
|---|---|---|---|---|---|
| Bin1001 | 97 | 6 | 17 | 93.1 | 5.98 |
| Bin1020 | 107 | 3 | 10 | 93.93 | 4.25 |
| Bin1036 | 84 | 6 | 30 | 91.67 | 7.77 |
| Bin1059 | 103 | 8 | 9 | 91.38 | 5.33 |
| Bin1091 | 101 | 9 | 10 | 95.68 | 2.25 |
| Bin1111 | 117 | 2 | 1 | 97.52 | 0.56 |
| Bin1152 | 117 | 1 | 2 | 96.24 | 0.54 |
| Bin1306 | 109 | 4 | 7 | 92.31 | 4.92 |
| Bin1359 | 107 | 2 | 11 | 91.28 | 0.87 |
| Bin1501 | 105 | 5 | 10 | 96.43 | 1.18 |
| Bin154 | 111 | 7 | 2 | 100 | 2.2 |
| Bin204 | 113 | 5 | 2 | 96.77 | 1.21 |
| Bin205 | 101 | 9 | 10 | 92.08 | 5.05 |
| Bin22 | 117 | 2 | 1 | 100 | 0.48 |
| Bin231 | 115 | 3 | 2 | 93.64 | 2 |
| Bin265 | 108 | 3 | 9 | 95.02 | 4.25 |
| Bin328 | 95 | 6 | 19 | 92.24 | 5.56 |
| Bin336 | 107 | 9 | 4 | 97.72 | 3.94 |
| Bin367 | 116 | 3 | 1 | 97.13 | 2.15 |
| Bin396 | 109 | 10 | 1 | 98.12 | 6.45 |
| Bin403 | 109 | 2 | 9 | 97.04 | 1.61 |
| Bin428 | 100 | 13 | 7 | 95.77 | 5.96 |
| Bin481 | 102 | 6 | 12 | 91.83 | 0.65 |
| Bin484 | 115 | 3 | 2 | 93.64 | 1.45 |
| Bin493 | 117 | 3 | 0 | 94.47 | 2 |

| Name | Unique Gene Count | Multiple Gene Count | Missing Gene Count | Genome completeness (%) | Contamination (%) |
|---|---|---|---|---|---|
| Bin50 | 106 | 13 | 1 | 97.85 | 3.46 |
| Bin583 | 114 | 1 | 5 | 98.9 | 1.1 |
| Bin609 | 105 | 3 | 12 | 95.1 | 4.3 |
| Bin617 | 108 | 10 | 2 | 91.81 | 5.18 |
| Bin631 | 104 | 8 | 8 | 96.63 | 4.49 |
| Bin684 | 106 | 4 | 10 | 90.34 | 2.03 |
| Bin737 | 114 | 0 | 6 | 93.96 | 1.46 |
| Bin744 | 98 | 8 | 14 | 91.39 | 2.32 |
| Bin768 | 110 | 3 | 7 | 95.48 | 6.3 |
| Bin785 | 99 | 15 | 6 | 91.43 | 5.12 |
| Bin790 | 93 | 7 | 20 | 90.34 | 3.53 |
| Bin803 | 118 | 1 | 1 | 94.57 | 0.55 |
| Bin820 | 112 | 4 | 4 | 94.84 | 0.22 |
| Bin931 | 110 | 2 | 8 | 95.76 | 1.15 |
| Bin967 | 111 | 2 | 7 | 99.26 | 0.84 |

### Table S5

*Table 5. Catabolite repression resistance genes (*phosphotransferase system sugar specific EII component *or* putative sugar kinases) *in good lipolytic MAGs.*

| MAGs ID | Gene quantity | Sugar | Gene name |
|---|---|---|---|
| 583 | 1 | PTS system fructose-specific EIIABC component | fruA_2 |
| | 1 | PTS system mannose-specific EIIAB component | manX, 2.7.1.191 |
| 803 | 1 | PTS system fructose-specific EIIABC component | fruA |
| 403 | 1 | putative sugar kinase YdjH | ydjH, 2.7.1.- |
| 396 | 1 | putative sugar kinase YdjH | ydjH, 2.7.1.- |
| 1152 | 2 | putative sugar kinase YdjH | ydjH, 2.7.1.- |
| 367 | 1 | putative sugar kinase YdjH | ydjH, 2.7.1.- |
| 50 | 1 | putative sugar kinase YdjH | ydjH, 2.7.1.- |
| 684 | 2 | PTS system fructose-specific EIIA component | fruA |
| 1036 | 2 | PTS system mannose-specific EIIAB component | manX, 2.7.1.191 |
| | 3 | putative sugar kinase YdjH ydjH, 2.7.1.- | ydjH, 2.7.1.- |
| 1091 | 1 | PTS system mannose-specific EIIAB component | manX, 2.7.1.191 |
| 1359 | 1 | PTS system mannose-specific EIIAB component | manX, 2.7.1.191 |
| 22 | 1 | PTS system mannose-specific EIIAB component | manX, 2.7.1.191 |
| 265 | 1 | PTS system mannose-specific EIIAB component | manX, 2.7.1.191 |
| 967 | 1 | PTS system mannose-specific EIIAB component | manX, 2.7.1.191 |
| | 1 | PTS system fructose-specific EIIB'BC component | fruA |
| 154 | 2 | PTS system fructose-specific EIIABC component | fruA |
| | 2 | putative sugar kinase YdjH | ydjH, 2.7.1.- |
| 609 | 1 | PTS system fructose-specific EIIABC component | fruA |
| 631 | 3 | PTS system fructose-specific EIIABC component | fruA |
| 820 | 1 | PTS system mannose-specific EIIBCA component | manP |
| | 1 | PTS system fructose-specific EIIABC component | fruA |
| | 2 | putative sugar kinase YdjH | ydjH, 2.7.1.- |
| 617 | 1 | putative sugar kinase YdjH | ydjH, 2.7.1.- |
| 1501 | 3 | PTS system fructose-specific EIIABC component | fruA |
| 481 | 0 | - | - |

| MAGs ID | Gene quantity | Sugar | Gene name |
|---|---|---|---|
| 484 | 2 | PTS system fructose-specific EIIB'BC component | fruA |
| | 1 | putative sugar kinase YdjH | ydjH, 2.7.1.- |
| 231 | 0 | - | - |
| 204 | 2 | putative sugar kinase YdjH | ydjH, 2.7.1.- |
| 1059 | 1 | putative sugar kinase YdjH | ydjH, 2.7.1.- |
| 1001 | 0 | - | - |
| 328 | 0 | - | - |
| 931 | 1 | putative sugar kinase YdjH | ydjH, 2.7.1.- |
| 336 | 1 | putative sugar kinase YdjH | ydjH, 2.7.1.- |
| 1020 | 1 | PTS system fructose-specific EIIABC component | fruA |
| | 1 | PTS system mannitol-specific EIICBA component | mtlA |
| | 1 | PTS system glucose-specific EIIA component | crr, 2.7.1.199 |
| | 1 | PTS system glucose-specific EIICBA component | ptsG, 2.7.1.199 |
| 744 | 1 | PTS system fructose-specific EIIABC component | fruA |
| | 1 | PTS system glucose-specific EIIA component | crr, 2.7.1.199 |
| | 1 | PTS system glucose-specific EIICBA component | ptsG, 2.7.1.199 |
| | 1 | PTS system beta-glucoside-specific EIIBCA component | bglF |
| 768 | 2 | PTS system glucose-specific EIICBA component | ptsG_1, 2.7.1.199 |
| | 1 | PTS system fructose-specific EIIABC component | fruA |
| 1111 | 1 | PTS system beta-glucoside-specific EIIBCA component | bglF |
| | 1 | PTS system fructose-specific EIIABC component | fruA |
| | 1 | PTS system glucose-specific EIIA component | crr, 2.7.1.199 |
| 493 | 1 | PTS system mannitol-specific EIICB component | mtlA |
| | 1 | PTS system beta-glucoside-specific EIIBCA component | bglF |
| | 1 | PTS system fructose-specific EIIABC component | fruA |
| 785 | 1 | PTS system beta-glucoside-specific EIIBCA component | bglF |
| 205 | 1 | putative sugar kinase YdjH | ydjH, 2.7.1.- |
| 790 | 0 | - | - |
| 1306 | 2 | PTS system fructose-specific EIIABC component | fruA |
| | 1 | PTS system mannitol-specific EIICBA component | mtlA |

| MAGs ID | Gene quantity | Sugar | Gene name |
|---|---|---|---|
| | 1 | PTS system beta-glucoside-specific EIIBCA component | bglF |
| | 1 | putative sugar kinase YdjH | ydjH_1, 2.7.1.- |
| 428 | 1 | PTS system beta-glucoside-specific EIIBCA component | bglF |
| | 1 | PTS system mannitol-specific EIICBA component | mtlA |
| | 2 | PTS system fructose-specific EIIB'BC component | fruA |
| | 1 | putative sugar kinase YdjH | ydjH, 2.7.1.- |
| 737 | 2 | Ascorbate-specific PTS system EIIB component | ulaB, 2.7.1.194 |
| | 2 | PTS system fructose-specific EIIABC component | fruA |
| | 1 | PTS system 2-O-alpha-mannosyl-D-glycerate-specific EIIABC component | mngA |

### Table S6

*Table 6. Grouping the putative lipolytic MAGs based on the role of the lipase on the genome.*

| MAG ID | Lowest classified level | Phyla | Gram stain | fadL | Denitrification/PHA genes |
|---|---|---|---|---|---|
| Bin1001.gff | Family- Obscuribacteraceae | Cyanobacteria | Gram negative | None | Only Denitrification |
| Bin1020.gff | Genus-Mycolicibacterium | Actinobacteriota | Gram positive | None | Only PHA |
| Bin1036.gff | Family-Anderseniellaceae | Proteobacteria | Gram negative | None | Both |
| Bin1059.gff | Family-Acutalibacteraceae | Firmicutes_A | Gram positive | None | None |
| Bin1091.gff | Genus-Paracoccus | Proteobacteria | Gram negative | None | Both |
| Bin1111.gff | Genus-Corynebacterium | Actinobacteriota | Gram positive | None | None |
| Bin1152.gff | Order-Bacteroidales | Bacteroidota | Gram negative | None | None |
| Bin1306.gff | Genus-Austwickia | Actinobacteriota | Gram positive | None | Only PHA |
| Bin1359.gff | Class-Gammaproteobacteria | Proteobacteria | Gram negative | None | None |
| Bin1501.gff | Class-Syntrophorhabdia | Desulfobacterota | Gram negative | Yes | None |
| Bin154.gff | Order-Hydrogenedentiales | Hydrogenedentota | Not known | None | Only Denitrification |
| Bin204.gff | Order-Christensenellales | Firmicutes_A | Gram positive | None | Only Denitrification |
| Bin205.gff | Order-Nanopelagicales | Actinobacteriota | Gram positive | None | Only PHA |
| Bin22.gff | Genus-Nitrosomonas | Proteobacteria | Gram negative | None | Only Denitrification |
| Bin231.gff | Order-Anaerolineales | Chloroflexota | Mostly gram negative | None | None |
| Bin265.gff | Family-Rhodocyclaceae | Proteobacteria | Gram negative | None | Only PHA |
| Bin328.gff | Family-Obscuribacteraceae | Cyanobacteria | Gram negative | None | None |
| Bin336.gff | Family-Microtrichaceae | Actinobacteriota | Gram positive | None | Only PHA |
| Bin367.gff | Genus-Lentimicrobium | Bacteroidota | Gram negative | None | Only Denitrification |
| Bin396.gff | Order-Flavobacteriales | Bacteroidota | Gram negative | None | Only Denitrification |
| Bin403.gff | Order-Flavobacteriales | Bacteroidota | Gram negative | None | Only Denitrification |
| Bin428.gff | Genus-Austwickia | Actinobacteriota | Gram positive | None | Both |
| Bin481.gff | Species-Desulfobacter postgatei | Desulfobacterota | Gram negative | None | None |
| Bin484.gff | Order-Anaerolineales | Chloroflexota | Mostly gram negative | None | Only Denitrification |
| Bin493.gff | Genus-Propionicimonas | Actinobacteriota | Gram positive | None | Only Denitrification |
| Bin50.gff | Order-Bacteroidales | Bacteroidota | Gram negative | None | Both |

| MAG ID | Lowest classified level | Phyla | Gram stain | fadL | Denitrification/PHA genes |
|---|---|---|---|---|---|
| Bin583.gff | Class-Krumholzibacteria | Krumholzibacteriota | Gram negative | None | None |
| Bin609.gff | Phylum-Omnitrophota | Omnitrophota | Not known | None | None |
| Bin617.gff | Class-Polyangia | Myxococcota | Gram negative | None | Only PHA |
| Bin631.gff | Phylum-Spirochaetota | Spirochaetota | Weak Gram negative in some | None | Only Denitrification |
| Bin684.gff | Unassigned | Unassigned | Not known | None | None |
| Bin737.gff | Genus-Rhodoluna | Actinobacteriota | Gram positive | None | None |
| Bin744.gff | Genus-Mycolicibacterium | Actinobacteriota | Gram positive | None | Only PHA |
| Bin768.gff | Genus-Mycolicibacterium | Actinobacteriota | Gram positive | None | Both |
| Bin785.gff | Genus-Propionicimonas | Actinobacteriota | Gram positive | None | Only Denitrification |
| Bin790.gff | Order-Nanopelagicales | Actinobacteriota | Gram positive | None | None |
| Bin803.gff | Genus-Chlorobium | Bacteroidota | Gram negative | None | None |
| Bin820.gff | Unassigned | Unassigned | Not known | None | None |
| Bin931.gff | Order-Solirubrobacterales | Actinobacteriota | Gram positive | None | None |
| Bin967.gff | Genus-Rhodoferax | Proteobacteria | Gram negative | Yes | Both |

### Table S7

*Table 7. Details of taxonomic classification for good lipolytic MAGs by GTDB-Tk.*

| User Genome | Classification | FastANI Reference[1] | FastANI Reference Radius[2] | FastANI Taxonomy[3] | FastANI ANI[4] | FastANI Alignment Fraction[5] | Closest Placement Reference[6] | Closest Placement Taxonomy[7] | Closest Placement ANI[8] | Closest Placement Alignment Fraction[9] | Classification Method[10] | AA Percent[11] | RED Value[12] |
|---|---|---|---|---|---|---|---|---|---|---|---|---|---|
| Bin 1001 | p__Cyanobacteria;<br>c__Vampirovibrionia;<br>o__Obscuribacterales;<br>f__Obscuribacteraceae;<br>g__Ga0077546;<br>s__ | | | | | | GCA_001464165.1 | p__Cyanobacteria;<br>c__Vampirovibrionia;<br>o__Obscuribacterales;<br>f__Obscuribacteraceae;<br>g__Ga0077546;<br>s__Ga0077546 sp001464165 | 84.39 | 0.67 | RED | 83.43 | 0.980 |
| Bin 1020 | p__Actinobacteriota;<br>c__Actinobacteria;<br>o__Mycobacteriales;<br>f__Mycobacteriaceae;<br>g__Mycolicibacterium;<br>s__ | | | | | | | | | | Topology | 89.38 | 0.963 |
| Bin 1036 | p__Proteobacteria;<br>c__Alphaproteobacteria;<br>o__Rhizobiales;<br>f__Anderseniellaceae;<br>g__QKVK01;<br>s__ | | | | | | GCF_003234965.1 | p__Proteobacteria;<br>c__Alphaproteobacteria;<br>o__Rhizobiales;<br>f__Anderseniellaceae;<br>g__QKVK01;<br>s__QKVK01 sp003234965 | 86.92 | 0.67 | RED | 72.48 | 0.982 |
| Bin 1059 | p__Firmicutes_A;<br>c__Clostridia;<br>o__Oscillospirales;<br>f__Acutalibacteraceae;<br>g__UBA1447;<br>s__ | | | | | | | | | | RED | 89.86 | 0.909 |
| Bin 1091 | p__Proteobacteria;<br>c__Alphaproteobacteria;<br>o__Rhodobacterales;<br>f__Rhodobacteraceae;<br>g__Paracoccus;<br>s__ | | | | | | | | | | RED | 89.86 | 0.944 |
| Bin 1111 | p__Actinobacteriota;<br>c__Actinobacteria;<br>o__Mycobacteriales;<br>f__Mycobacteriaceae;<br>g__Corynebacterium;<br>s__ | | | | | | | | | | Topology | 97 | 0.986 |

| User Genome | Classification | FastANI Reference[1] | FastANI Reference Radius[2] | FastANI Taxonomy[3] | FastANI ANI[4] | FastANI Alignment Fraction[5] | Closest Placement Reference[6] | Closest Placement Taxonomy[7] | Closest Placement ANI[8] | Closest Placement Alignment Fraction[9] | Classification Method[10] | AA Percent[11] | RED Value[12] |
|---|---|---|---|---|---|---|---|---|---|---|---|---|---|
| Bin 1152 | p__Bacteroidota;<br>c__Bacteroidia;<br>o__Bacteroidales;<br>f__WCHB1-69;<br>g__UBA5266;<br>s__ | | | | | | GCA_002411 545.1 | p__Bacteroidota;<br>c__Bacteroidia;<br>o__Bacteroidales;<br>f__WCHB1-69;<br>g__UBA5266;<br>s__UBA5266 sp002411545 | 77.31 | 0.2 | RED | 95.22 | 0.923 |
| Bin 1306 | p__Actinobacteriota;<br>c__Actinobacteria;<br>o__Actinomycetales;<br>f__Dermatophilaceae;<br>g__Austwickia;<br>s__ | | | | | | GCF_000298 175.1 | p__Actinobacteriota;<br>c__Actinobacteria;<br>o__Actinomycetales;<br>f__Dermatophilaceae;<br>g__Austwickia;<br>s__Austwickia chelonae | 78.24 | 0.27 | RED | 90.97 | 0.898 |
| Bin 1359 | p__Proteobacteria;<br>c__Gammaproteobacteria;<br>o__UBA6002;<br>f__UBA6002;<br>g__;<br>s__ | | | | | | | | | | RED | 90.04 | 0.745 |
| Bin 1501 | p__Desulfobacterota;<br>c__Syntrophorhabdia;<br>o__;<br>f__;<br>g__;<br>s__ | | | | | | | | | | RED | 89.11 | 0.434 |
| Bin 154 | p__Hydrogenedentota;<br>c__Hydrogenedentia;<br>o__Hydrogenedentiales;<br>f__;<br>g__;<br>s__ | | | | | | | | | | RED | 96.15 | 0.715 |
| Bin 204 | p__Firmicutes_A;<br>c__Clostridia;<br>o__Christensenellales;<br>f__CAG-74;<br>g__DTU024;<br>s__ | | | | | | GCA_002428 405.1 | p__Firmicutes_A;<br>c__Clostridia;<br>o__Christensenellales;<br>f__CAG-74;<br>g__DTU024;<br>s__DTU024 sp002428405 | 77.63 | 0.17 | Topology | 95.36 | 0.946 |

| User Genome | Classification | FastANI Reference[1] | FastANI Reference Radius[2] | FastANI Taxonomy[3] | FastANI ANI[4] | FastANI Alignment Fraction[5] | Closest Placement Reference[6] | Closest Placement Taxonomy[7] | Closest Placement ANI[8] | Closest Placement Alignment Fraction[9] | Classification Method[10] | AA Percent[11] | RED Value[12] |
|---|---|---|---|---|---|---|---|---|---|---|---|---|---|
| Bin 205 | d__Bacteria; p__Actinobacteriota; c__Actinobacteria; o__Nanopelagicales; f__GCA-2699445; g__; s__ | | | | | | | | | | RED | 89.03 | 0.770 |
| Bin 22 | p__Proteobacteria; c__Gammaproteobacteria; o__Burkholderiales; f__Nitrosomonadaceae; g__Nitrosomonas; s__ | | | | | | GCF_003201565.1 | p__Proteobacteria; c__Gammaproteobacteria; o__Burkholderiales; f__Nitrosomonadaceae; g__Nitrosomonas; s__Nitrosomonas sp003201565 | 77.59 | 0.25 | Topology | 98.25 | 0.953 |
| Bin 231 | p__Chloroflexota; c__Anaerolineae; o__Anaerolineales; f__envOPS12; g__; s__ | | | | | | | | | | Topology | 95.4 | 0.903 |
| Bin 265 | p__Proteobacteria; c__Gammaproteobacteria; o__Burkholderiales; f__Rhodocyclaceae; g__; s__ | | | | | | | | | | RED | 91.47 | 0.916 |
| Bin 328 | p__Cyanobacteria; c__Vampirovibrionia; o__Obscuribacterales; f__Obscuribacteraceae; g__Ga0077546; s__ | | | | | | GCA_001464165.1 | p__Cyanobacteria; c__Vampirovibrionia; o__Obscuribacterales; f__Obscuribacteraceae; g__Ga0077546; s__Ga0077546 sp001464165 | 86.62 | 0.74 | RED | 81.39 | 0.985 |
| Bin 336 | p__Actinobacteriota; c__Acidimicrobiia; o__Microtrichales; f__Microtrichaceae; g__IMCC26207; s__ | | | | | | GCF_001025035.1 | p__Actinobacteriota; c__Acidimicrobiia; o__Microtrichales; f__Microtrichaceae; g__IMCC26207; s__IMCC26207 sp001025035 | 76.31 | 0.05 | RED | 93.71 | 0.857 |

| User Genome | Classification | FastANI Reference[1] | FastANI Reference Radius[2] | FastANI Taxonomy[3] | FastANI ANI[4] | FastANI Alignment Fraction[5] | Closest Placement Reference[6] | Closest Placement Taxonomy[7] | Closest Placement ANI[8] | Closest Placement Alignment Fraction[9] | Classification Method[10] | AA Percent[11] | RED Value[12] |
|---|---|---|---|---|---|---|---|---|---|---|---|---|---|
| Bin 367 | p__Bacteroidota;<br>c__Bacteroidia;<br>o__Bacteroidales;<br>f__Lentimicrobiaceae;<br>g__Lentimicrobium;<br>s__ | | | | | | GCA_002426 025.1 | p__Bacteroidota;<br>c__Bacteroidia;<br>o__Bacteroidales;<br>f__Lentimicrobiaceae;<br>g__Lentimicrobium;<br>s__Lentimicrobium sp002426025 | 77.09 | 0.17 | Topology | 97.48 | 0.949 |
| Bin 396 | p__Bacteroidota;<br>c__Bacteroidia;<br>o__Flavobacteriales;<br>f__PHOS-HE28;<br>g__PHOS-HE28;<br>s__ | | | | | | GCA_002342 985.1 | p__Bacteroidota;<br>c__Bacteroidia;<br>o__Flavobacteriales;<br>f__PHOS-HE28;<br>g__PHOS-HE28;<br>s__PHOS-HE28 sp002342985 | 82.57 | 0.55 | Topology | 97.22 | 0.943 |
| Bin 403 | p__Bacteroidota;<br>c__Bacteroidia;<br>o__Flavobacteriales;<br>f__PHOS-HE28;<br>g__PHOS-HE28;<br>s__ | | | | | | GCA_002396 605.1 | p__Bacteroidota;<br>c__Bacteroidia;<br>o__Flavobacteriales;<br>f__PHOS-HE28;<br>g__PHOS-HE28;<br>s__PHOS-HE28 sp002396605 | 79.16 | 0.47 | Topology | 90.44 | 0.949 |
| Bin 428 | p__Actinobacteriota;<br>c__Actinobacteria;<br>o__Actinomycetales;<br>f__Dermatophilaceae;<br>g__Austwickia;<br>s__ | | | | | | GCF_000298 175.1 | p__Actinobacteriota;<br>c__Actinobacteria;<br>o__Actinomycetales;<br>f__Dermatophilaceae;<br>g__Austwickia;<br>s__Austwickia chelonae | 78.86 | 0.28 | RED | 91.15 | 0.896 |
| Bin 481 | p__Desulfobacterota;<br>c__Desulfobacteria;<br>o__Desulfobacterales;<br>f__Desulfobacteraceae;<br>g__Desulfobacter;<br>s__Desulfobacter postgatei | GCF_000233 695.2 | 95 | p__Desulfobacterota;<br>c__Desulfobacteria;<br>o__Desulfobacterales;<br>f__Desulfobacteraceae;<br>g__Desulfobacter;<br>s__Desulfobacter postgatei | 96.18 | 0.81 | GCF_000233 695.2 | p__Desulfobacterota;<br>c__Desulfobacteria;<br>o__Desulfobacterales;<br>f__Desulfobacteraceae;<br>g__Desulfobacter;<br>s__Desulfobacter postgatei | 96.18 | 0.81 | Topology and ANI | 87.52 | |
| Bin 484 | p__Chloroflexota;<br>c__Anaerolineae;<br>o__Anaerolineales;<br>f__envOPS12;<br>g__;<br>s__ | | | | | | | | | | Topology | 95.26 | 0.903 |

| User Genome | Classification | FastANI Reference[1] | FastANI Reference Radius[2] | FastANI Taxonomy[3] | FastANI ANI[4] | FastANI Alignment Fraction[5] | Closest Placement Reference[6] | Closest Placement Taxonomy[7] | Closest Placement ANI[8] | Closest Placement Alignment Fraction[9] | Classification Method[10] | AA Percent[11] | RED Value[12] |
|---|---|---|---|---|---|---|---|---|---|---|---|---|---|
| Bin 493 | p__Actinobacteriota; c__Actinobacteria; o__Propionibacteriales; f__Propionibacteriaceae; g__Propionicimonas; s__ | | | | | | GCA_002841335.1 | p__Actinobacteriota; c__Actinobacteria; o__Propionibacteriales; f__Propionibacteriaceae; g__Propionicimonas; s__Propionicimonas sp002841335 | 86.39 | 0.84 | Topology | 97.02 | 0.987 |
| Bin 50 | p__Bacteroidota; c__Bacteroidia; o__Bacteroidales; f__4484-276; g__; s__ | | | | | | | | | | RED | 96.15 | 0.800 |
| Bin 583 | p__Krumholzibacteriota; c__Krumholzibacteria; o__SSS58A; f__SSS58A; g__; s__ | | | | | | | | | | RED | 94.05 | 0.856 |
| Bin 609 | p__Omnitrophota; c__koll11; o__UBA1560; f__2-01-FULL-45-10; g__FEN-1322; s__ | | | | | | GCA_003140915.1 | p__Omnitrophota; c__koll11; o__UBA1560; f__2-01-FULL-45-10; g__FEN-1322; s__FEN-1322 sp003140915 | 76.57 | 0.21 | RED | 87.16 | 0.910 |
| Bin 617 | p__Myxococcota; c__Polyangia; o__HGW-17; f__; g__; s__ | | | | | | | | | | RED | 95.85 | 0.588 |
| Bin 631 | p__Spirochaetota; c__UBA4802; o__UBA4802; f__UBA5368; g__; s__ | | | | | | GCA_002407865.1 | p__Spirochaetota; c__UBA4802; o__UBA4802; f__UBA5368; g__UBA5368; s__UBA5368 sp002407865 | 76.69 | 0.11 | RED | 90.38 | 0.811 |

| User Genome | Classification | FastANI Reference[1] | FastANI Reference Radius[2] | FastANI Taxonomy[3] | FastANI ANI[4] | FastANI Alignment Fraction[5] | Closest Placement Reference[6] | Closest Placement Taxonomy[7] | Closest Placement ANI[8] | Closest Placement Alignment Fraction[9] | Classification Method[10] | AA Percent[11] | RED Value[12] |
|---|---|---|---|---|---|---|---|---|---|---|---|---|---|
| Bin 684 | p__UBA10199; c__UBA10199; o__GCA-002796325; f__1-14-0-20-49-13; g__; s__ | | | | | | | | | | RED | 87.84 | 0.783 |
| Bin 737 | p__Actinobacteriota; c__Actinobacteria; o__Actinomycetales; f__Microbacteriaceae; g__Rhodoluna; s__ | | | | | | GCF_000699505.1 | p__Actinobacteriota; c__Actinobacteria; o__Actinomycetales; f__Microbacteriaceae; g__Rhodoluna; s__Rhodoluna lacicola | 79.25 | 0.41 | Topology | 92.86 | 0.952 |
| Bin 744 | p__Actinobacteriota; c__Actinobacteria; o__Mycobacteriales; f__Mycobacteriaceae; g__Mycolicibacterium; s__ | | | | | | GCA_001510415.1 | p__Actinobacteriota; c__Actinobacteria; o__Mycobacteriales; f__Mycobacteriaceae; g__Mycolicibacterium; s__Mycolicibacterium sp001510415 | 80.57 | 0.59 | Topology | 85.91 | 0.967 |
| Bin 768 | p__Actinobacteriota; c__Actinobacteria; o__Mycobacteriales; f__Mycobacteriaceae; g__Mycolicibacterium; s__ | | | | | | GCA_001510415.1 | p__Actinobacteriota; c__Actinobacteria; o__Mycobacteriales; f__Mycobacteriaceae; g__Mycolicibacterium; s__Mycolicibacterium sp001510415 | 77.91 | 0.33 | Topology | 91.31 | 0.956 |
| Bin 785 | p__Actinobacteriota; c__Actinobacteria; o__Propionibacteriales; f__Propionibacteriaceae; g__Propionicimonas; s__ | | | | | | | | | | Topology | 91.69 | 0.983 |
| Bin 790 | p__Actinobacteriota; c__Actinobacteria; o__Nanopelagicales; f__UBA10799; g__UBA10799; s__ | | | | | | GCA_003452655.1 | p__Actinobacteriota; c__Actinobacteria; o__Nanopelagicales; f__UBA10799; g__UBA10799; s__UBA10799 sp003452655 | 78.04 | 0.1 | RED | 80.97 | 0.861 |

| User Genome | Classification | FastANI Reference[1] | FastANI Reference Radius[2] | FastANI Taxonomy[3] | FastANI ANI[4] | FastANI Alignment Fraction[5] | Closest Placement Reference[6] | Closest Placement Taxonomy[7] | Closest Placement ANI[8] | Closest Placement Alignment Fraction[9] | Classification Method[10] | AA Percent[11] | RED Value[12] |
|---|---|---|---|---|---|---|---|---|---|---|---|---|---|
| Bin 803 | p__Bacteroidota; c__Chlorobia; o__Chlorobiales; f__Chlorobiaceae; g__Chlorobium; s__ | | | | | | | | | | Topology | 97.02 | 0.944 |
| Bin 820 | p__RBG-13-61-14; c__RBG-13-61-14; o__RBG-13-61-14; f__; g__; s__ | | | | | | GCA_001797815.1 | p__RBG-13-61-14; c__RBG-13-61-14; o__RBG-13-61-14; f__RBG-13-61-14; g__RBG-13-61-14; s__RBG-13-61-14 sp001797815 | 76.29 | 0.16 | RED | 95.58 | 0.639 |
| Bin 931 | p__Actinobacteriota; c__Thermoleophilia; o__Solirubrobacterales; f__70-9; g__67-14; s__ | | | | | | GCA_001897355.1 | p__Actinobacteriota; c__Thermoleophilia; o__Solirubrobacterales; f__70-9; g__67-14; s__67-14 sp001897355 | 82.74 | 0.7 | RED | 89.98 | 0.968 |
| Bin 967 | p__Proteobacteria; c__Gammaproteobacteria; o__Burkholderiales; f__Burkholderiaceae; g__Rhodoferax; s__ | | | | | | | | | | Topology | 93.08 | 0.981 |

1. Indicates the accession number of the closest reference genome as determine by ANI. This genome is used along with the placement of the genome in the reference tree to determine the species assignment on the genome. ANI values are only calculated when a query genome is placed within a defined genus and are calculated for all reference genomes in the genus 2. indicates the species-specific ANI circumscription radius of the reference genomes used to determine if a query genome should be classified to the same species as the reference 3. Indicates the GTDB taxonomy of the closest reference genome 4. Indicates the ANI between the query and the closest reference genome 5. Indicates the AF between the query and the closest reference genome 6. Indicates the accession number of the reference genome when a genome is placed on a terminal branch. This genome is used along with the ANI information to determine the species assignment on the genome 7. Indicates the GTDB taxonomy of the reference genome 8. Indicates the ANI between the query and the reference genome 9. Indicates the AF between the query and the reference genome 10. Indicates the rule used to classify the genome. This field will be one of: i) ANI/Placement, indicating a species assignment was made based on both the calculate ANI and placement of the genome in the reference tree; ii) taxonomic classification fully defined by topology, indicating that the classification could be determine based solely on the genome's position in the reference tree; or iii) taxonomic novelty determined using RED, indicating that the relative evolutionary divergence (RED) and placement of the genome in the reference tree were used to determine the classification 11. Indicates the percentage of the MSA spanned by the genome (i.e. percentage of columns with an amino acid) 12. Indicates, when required, the relative evolutionary divergence (RED) for a query genome. RED is not calculated when a query genome can be classified based on ANI.

#### Table S8

*Table 8. P-values from a two-way ANOVA on MAGs and reactor conditions including phase, treatment and temperature (α=0.05); highlighted cells in yellow had P-value≤0.05.*

| MAG_ID | P-value | | | MAG_ID | P-value | | |
|---|---|---|---|---|---|---|---|
| | Phase | Treatment | Temperature | | Phase | Treatment | Temperature |
| Bin 1001 | 0.037 | 0.604 | 0.134 | Bin 403 | 0.881 | 0.329 | 0.925 |
| Bin 1020 | 0.653 | 0.865 | 0.886 | Bin 428 | 0.255 | 0.045 | 0.597 |
| Bin 1036 | 0.548 | 0.797 | 0.768 | Bin 481 | 0.032 | 0.451 | 0.084 |
| Bin 1059 | 0.654 | 0.224 | 0.142 | Bin 484 | 0.343 | 0.082 | 0.16 |
| Bin 1091 | 0.672 | 0.587 | 0.842 | Bin 493 | 0.563 | 0.627 | 0.734 |
| Bin 1111 | 0.409 | 0.543 | 0.972 | Bin 50 | 0.807 | 0.968 | 0.211 |
| Bin 1152 | 0.216 | 0.807 | 0.101 | Bin 583 | 0.742 | 0.674 | 0.364 |
| Bin 1306 | 0.449 | 0.184 | 0.674 | Bin 609 | 0.006 | 0.267 | 0.001 |
| Bin 1359 | 0.846 | 0.559 | 0.957 | Bin 617 | 0.575 | 0.625 | 0.316 |
| Bin 1501 | 0.861 | 0.78 | 0.816 | Bin 631 | 0.852 | 0.26 | 0.179 |
| Bin 154 | 0.418 | 0.996 | 0.03 | Bin 684 | 0.87 | 0.753 | 0.664 |
| Bin 204 | 0.421 | 0.746 | 0.07 | Bin 737 | 0.547 | 0.387 | 0.899 |
| Bin 205 | 0.286 | 0.08 | 0.987 | Bin 744 | 0.665 | 0.926 | 0.763 |
| Bin 22 | 0.375 | 0.004 | 0.595 | Bin 768 | 0.671 | 0.912 | 0.864 |
| Bin 231 | 0.277 | 0.025 | 0.021 | Bin 785 | 0.762 | 0.406 | 0.929 |
| Bin 265 | 0.412 | 0.327 | 0.894 | Bin 790 | 0 | 0.129 | 0.798 |
| Bin 328 | 0 | 0.059 | 0.045 | Bin 803 | 0.352 | 0 | 0 |
| Bin 336 | 0.621 | 0.86 | 0.929 | Bin 820 | 0.85 | 0.824 | 0.101 |
| Bin 367 | 0.902 | 0.022 | 0.641 | Bin 931 | 0.72 | 0.649 | 0.897 |
| Bin 396 | 0.787 | 0.391 | 0.868 | | | | |

#### Figure S1

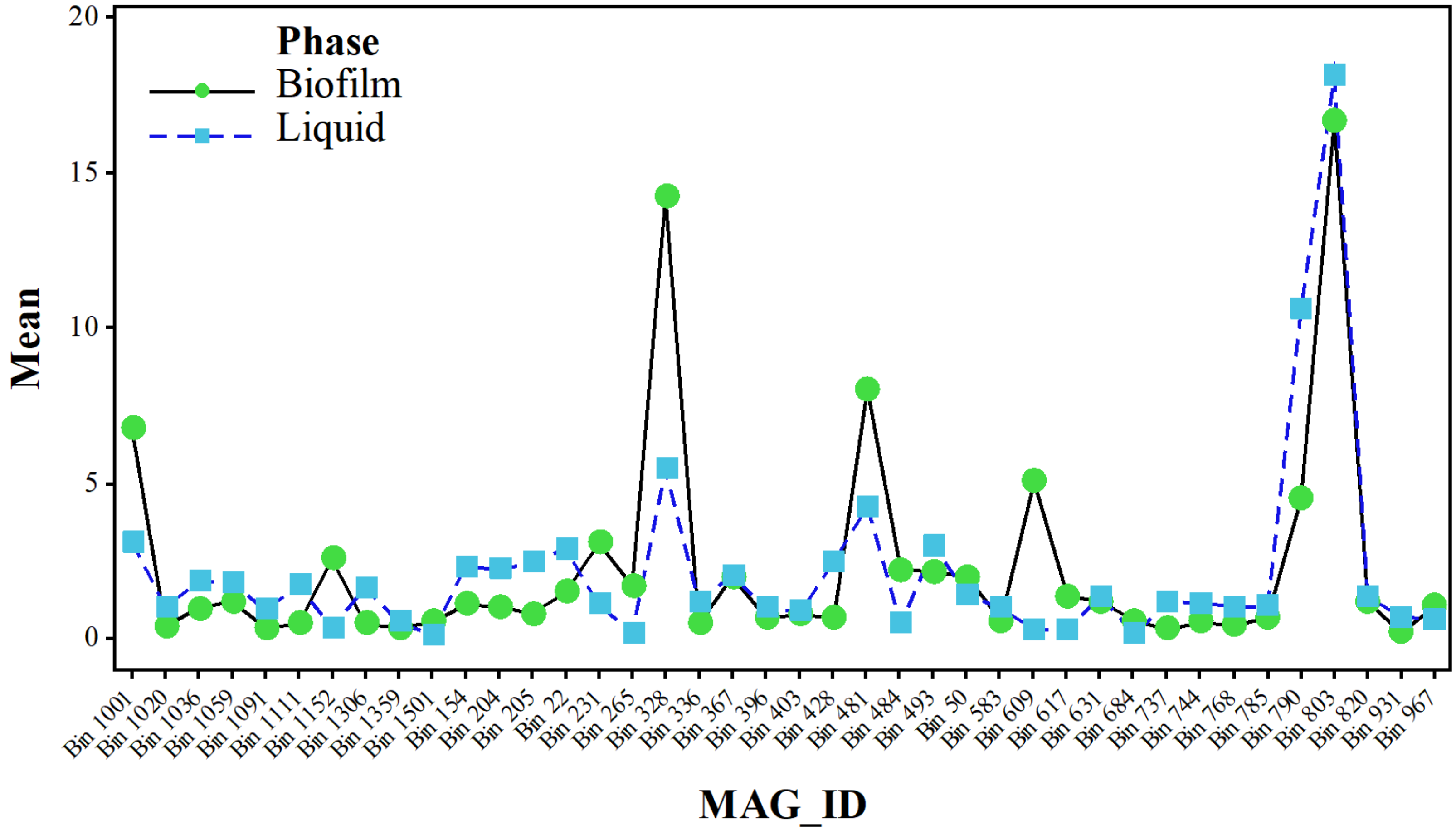


*Figure 1. Two-way ANOVA interaction plot (Minitab 18) for the abundance of reads per MAGs mapped to different phases (Biofilm and bulk Liquid) in the reactors.*

#### Figure S2

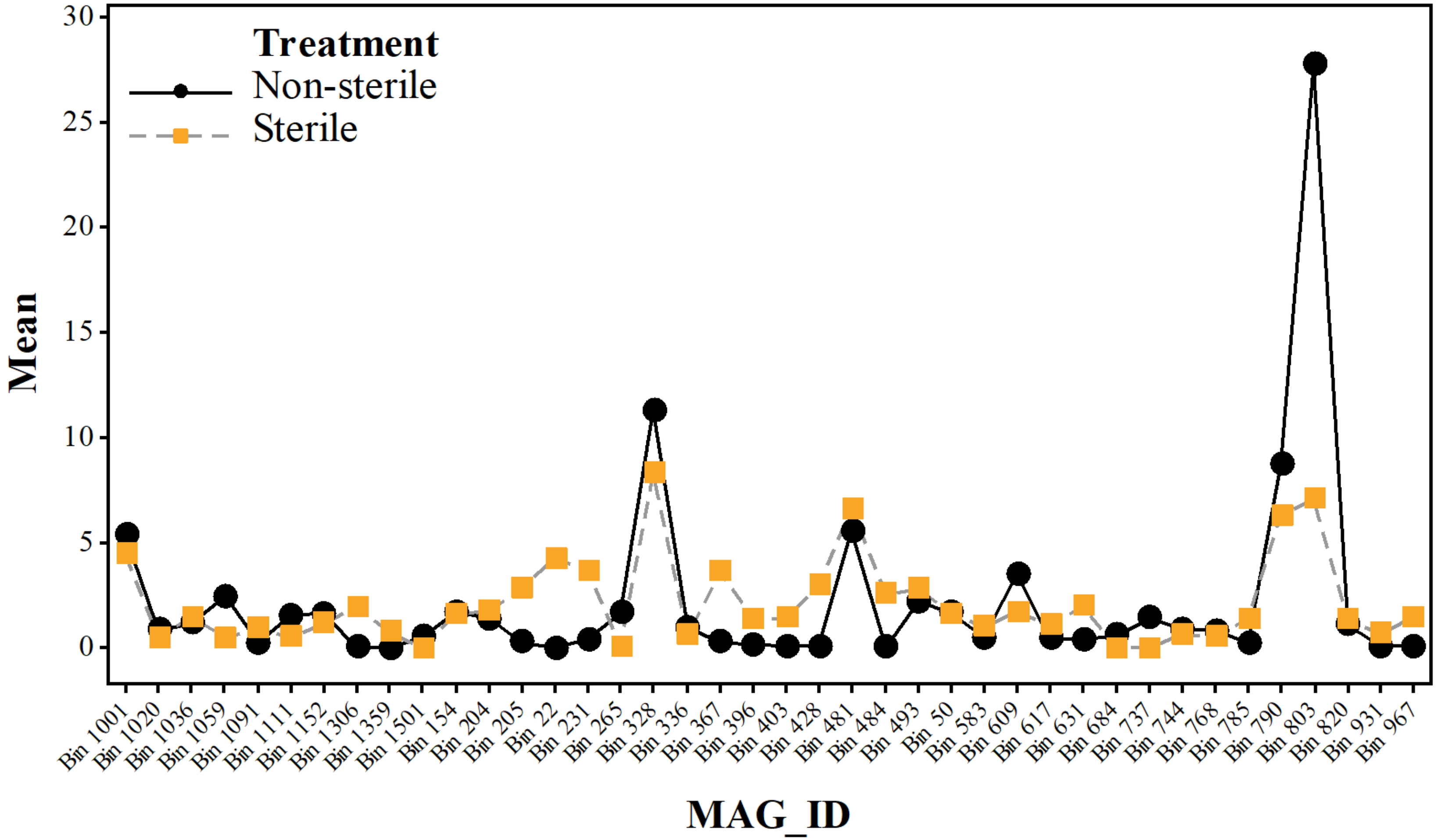


*Figure 2. Two-way ANOVA interaction plot (Minitab 18) for the abundance of reads per MAGs mapped to different treatment (Sterile and Non-sterile) in the reactors.*

#### Figure S3

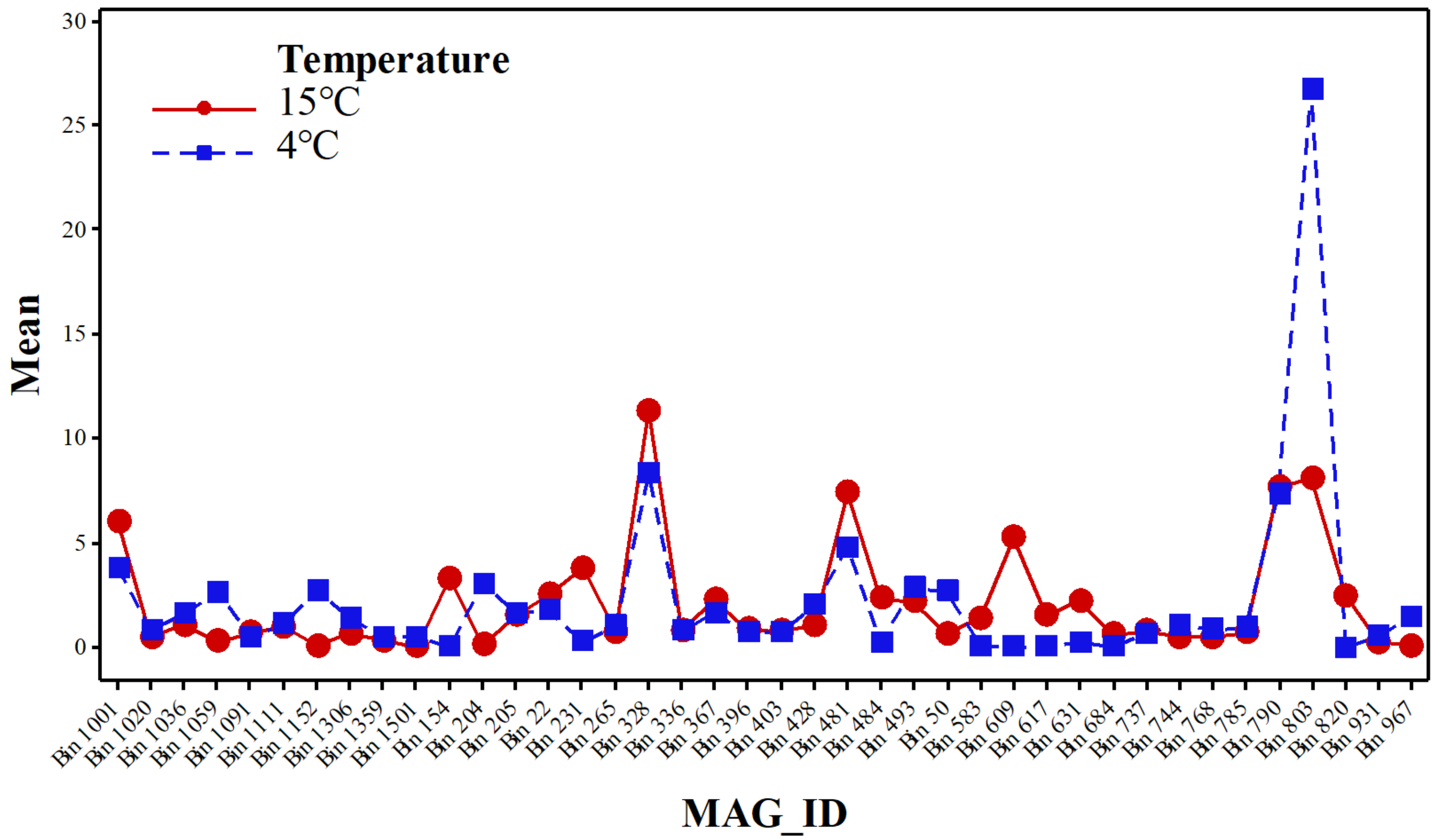


*Figure 3. Two-way ANOVA interaction plot (Minitab 18) for the abundance of reads per MAGs mapped to different temperature (4℃ and 15℃) in the reactors.*

### Table S9

*Table 9. MAGs linked to the taxa, reactor conditions and lipases, the status of the conditions in each MAG are based on the ANOVA analysis in Table 8.*

| MAGs ID | Cat.[1] | Phylum | Lowest classified level | | Treatment | Phase | Temp. (°C) | Lip. Quant[2] | Class | Length (aa) |
|---|---|---|---|---|---|---|---|---|---|---|
| 583 | 1 | Krumholzibacteriota | Class | Krumholzibacteria | Ster[3] ~ Nster[4] | Liq[5] ~ Bio[6] | 4 > 15 | 1 | Lipase 2 | 274 |
| 803 | 3 | Bacteroidota | Genus | Chlorobium | Nster >> Ster | Liq > Bio | 4 >> 15 | 1 | Lipase 1 | 287 |
| 403 | 1 | Bacteroidota | Order | Flavobacteriales | Ster > Nster | Liq ~ Bio | 4 ~ 15 | 1 | Lipase 3 | 362 |
| 396 | 6 | Bacteroidota | Order | Flavobacteriales | Ster > Nster | Liq ~ Bio | 4 ~ 15 | 1 | Lipase 3 | 363 |
| 1152 | 1 | Bacteroidota | Order | Bacteroidales | Ster ~ Nster | Bio > Liq | 4 > 15 | 1 | Lipase 2 | 321 |
| 367 | 2 | Bacteroidota | Genus | Lentimicrobium | Ster >> Nster | Liq ~ Bio | 4 ~ 15 | 1 | Lipase 2 | 305 |
| 50 | 2 | Bacteroidota | Order | Bacteroidales | Ster ~ Nster | Liq ~ Bio | 4 > 15 | 1 | Lipase 1 | 265 |
| 684 | 5 | Unassigned | - | - | Ster ~ Nster | Liq ~ Bio | 4 ~ 15 | 1 | Lipase | 247 |
| 1036 | 4 | Proteobacteria | Family | Anderseniellaceae | Ster ~ Nster | Liq > Bio | 4 ~ 15 | 1 | Putative | 391 |
| 1091 | 2 | Proteobacteria | Genus | Paracoccus | Ster ~ Nster | Liq ~ Bio | 4 ~ 15 | 1 | Lipase 3 | 294 |
| 1359 | 3 | Proteobacteria | Class | Gammaproteobacteria | Ster ~ Nster | Liq ~ Bio | 4 ~ 15 | 1 | Est A | 215 |
| 22 | 1 | Proteobacteria | Genus | Nitrosomonas | Ster >> Nster | Liq > Bio | 4 ~ 15 | 1 | Lipase 3 | 320 |
| 265 | 2 | Proteobacteria | Family | Rhodocyclaceae | Nster > Ster | Bio > Liq | 4 ~ 15 | 1 | Lipase 1 | 325 |
| 967 | 1 | Proteobacteria | Genus | Rhodoferax | Ster > Nster | Liq ~ Bio | 4 > 15 | 1 | Lipase | 306 |
| 154 | 2 | Hydrogenedentota | Order | Hydrogenedentiales | Ster ~ Nster | Liq > Bio | 15 >> 4 | 1 | Lipase 2 | 306 |
| 609 | 2 | Omnitrophota | Phylum | Omnitrophota | Nster > Ster | Bio >> Liq | 15 >> 4 | 1 | Lipase 1 | 220 |
| 631 | 2 | Spirochaetota | Phylum | Spirochaetota | Ster > Nster | Liq ~ Bio | 15 > 4 | 1 | Lactonizing | 297 |
| 820 | 3 | Unassigned | - | - | Ster ~ Nster | Liq ~ Bio | 15 > 4 | 1 | Lipase 2 | 308 |
| 617 | 4 | Myxococcota | Class | Polyangia | Ster ~ Nster | Bio > Liq | 15 > 4 | 4 | Lipase | 423 |
| | | | | | | | | | | 442 |
| | | | | | | | | | Est A | 289 |
| | | | | | | | | | Lipase 2 | 419 |
| 1501 | 1 | Desulfobacterota | Class | Syntrophorhabdia | Ster ~ Nster | Liq ~ Bio | 4 ~ 15 | 1 | Lactonizing | 253 |
| 481 | | Desulfobacterota | Species | Desulfobacter postgatei | Ster > Nster | Bio >> Liq | 15 >> 4 | 1 | Lipase 3 | 312 |
| 484 | 3 | Chloroflexota | Order | Anaerolineales | Ster > Nster | Bio > Liq | 15 > 4 | 3 | Lipase 1 | 274 |
| | | | | | | | | | | 246 |
| | | | | | | | | | Est A | 618 |
| 231 | 3 | Chloroflexota | Order | Anaerolineales | Ster >> Nster | Bio > Liq | 15 >> 4 | 1 | Lipase 3 | 246 |
| 204 | 1 | Firmicutes_A | Order | Christensenellales | Ster ~ Nster | Liq > Bio | 4 > 15 | 1 | Lipase 1 | 339 |

| MAGs ID | Cat.[1] | Phylum | Lowest classified level | | Treatment | Phase | Temp. (°C) | Lip. Quant[2] | Class | Length (aa) |
|---|---|---|---|---|---|---|---|---|---|---|
| 1059 | 4 | Firmicutes_A | Family | Acutalibacteraceae | Nster > Ster | Liq ~ Bio | 4 > 15 | 3 | Lipase | 211 |
| | | | | | | | | | | 561 |
| | | | | | | | | | | 404 |
| 1001 | 4 | Cyanobacteria | Family | Obscuribacteraceae | Ster ~ Nster | Bio >> Liq | 15 > 4 | 3 | Lipase 2 | 293 |
| | | | | | | | | | | 313 |
| | | | | | | | | | | 333 |
| 328 | 4 | Cyanobacteria | Family | Obscuribacteraceae | Nster >> Ster | Bio >> Liq | 15 >> 4 | 1 | Lipase 2 | 317 |
| 931 | 1 | Actinobacteriota | Order | Solirubrobacterales | Ster ~ Nster | Liq ~ Bio | 4 ~ 15 | 2 | Lipase 3 | 264 |
| | | | | | | | | | Putative | 414 |
| 336 | 2 | Actinobacteriota | Family | Microtrichaceae | Ster ~ Nster | Liq ~ Bio | 4 ~ 15 | 2 | Lipase | 319 |
| | | | | | | | | | Putative | 180 |
| 1020 | 5 | Actinobacteriota | Genus | Mycolicibacterium | Ster ~ Nster | Liq ~ Bio | 4 ~ 15 | 6 | Lipase 2 | 397 |
| | | | | | | | | | | 362 |
| | | | | | | | | | Triacylglycerol | 562 |
| | | | | | | | | | | 570 |
| | | | | | | | | | Putative | 445 |
| | | | | | | | | | | 412 |
| 744 | 5 | Actinobacteriota | Genus | Mycolicibacterium | Ster ~ Nster | Liq ~ Bio | 4 ~ 15 | 2 | Lipase | 348 |
| | | | | | | | | | Putative | 446 |
| 768 | 6 | Actinobacteriota | Genus | Mycolicibacterium | Ster ~ Nster | Liq ~ Bio | 4 ~ 15 | 7 | Lipase | 352 |
| | | | | | | | | | | 353 |
| | | | | | | | | | Lipase 2 | 291 |
| | | | | | | | | | | 253 |
| | | | | | | | | | Putative | 445 |
| | | | | | | | | | Triacylglycerol | 477 |
| | | | | | | | | | | 537 |
| 1111 | 1 | Actinobacteriota | Genus | Corynebacterium | Ster ~ Nster | Liq > Bio | 4 ~15 | 7 | Lipase | 339 |
| | | | | | | | | | Lipase 2 | 250 |
| | | | | | | | | | | 273 |
| | | | | | | | | | | 298 |
| | | | | | | | | | Putative | 450 |
| | | | | | | | | | | 456 |
| | | | | | | | | | | 471 |

| MAGs ID | Cat.[1] | Phylum | Lowest classified level | | Treatment | Phase | Temp. (°C) | Lip. Quant[2] | Class | Length (aa) |
|---|---|---|---|---|---|---|---|---|---|---|
| 493 | 3 | Actinobacteriota | Genus | Propionicimonas | Ster ~ Nster | Liq > Bio | 4 ~ 15 | 3 | Lipase 2 | 258 |
| | | | | | | | | | Lipase 3 | 720 |
| | | | | | | | | | Putative | 565 |
| 785 | 4 | Actinobacteriota | Genus | Propionicimonas | Ster > Nster | Liq ~ Bio | 4 ~ 15 | 1 | Putative | 569 |
| 205 | 4 | Actinobacteriota | Order | Nanopelagicales | Ster > Nster | Liq > Bio | 4 ~ 15 | 1 | Lipase 1 | 293 |
| 790 | 5 | Actinobacteriota | Order | Nanopelagicales | Nster > Ster | Liq >> Bio | 4 ~ 15 | 4 | Lipase 1 | 361 |
| | | | | | | | | | Lipase 3 | 308 |
| | | | | | | | | | Putative | 420 |
| | | | | | | | | | | 443 |
| 1306 | 5 | Actinobacteriota | Genus | Austwickia | Ster > Nster | Liq > Bio | 4 ~ 15 | 3 | Lipase 1 | 358 |
| | | | | | | | | | Triacylglycerol | 306 |
| | | | | | | | | | Putative | 368 |
| 428 | 6 | Actinobacteriota | Genus | Austwickia | Ster >> Nster | Liq > Bio | 4 ~ 15 | 3 | Lipase | 819 |
| | | | | | | | | | Lipase 1 | 339 |
| | | | | | | | | | Putative | 369 |
| 737 | 3 | Actinobacteriota | Genus | Rhodoluna | Nster > Ster | Liq > Bio | 4 ~ 15 | 1 | Lipase 3 | 311 |

1- Category 2- Lipase quantity 3- Sterile 4- Non-sterile 5- Liquid 6- Biofilm

#### Figure S4

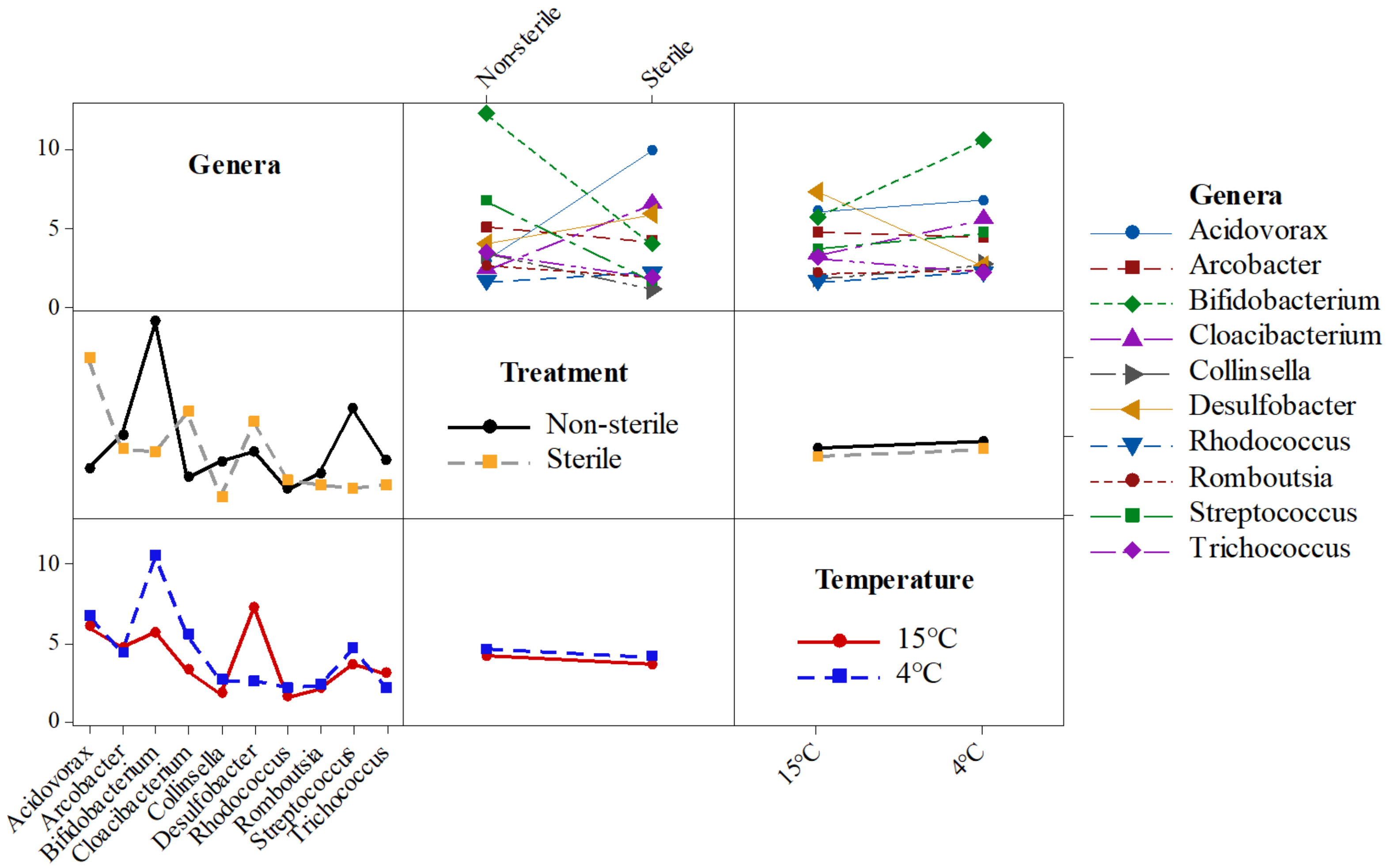


*Figure 4. Interaction plot (ANOVA, Minitab 18): Effect of temperature (4°C and 15°C) and treatment (sterile and non-sterile) on relative abundance of microbes at genus level.*

#### Table S10

*Table 10. List of identified proteins at FDR 1% and 5% by PEAKS two-round search.*

| Protein name | Gene name | FDR 1% | FDR 5% |
|---|---|---|---|
| Outer membrane porin protein 32 | omp32 | 73 | 81 |
| Vitamin B12 transporter BtuB | btuB | 14 | 15 |
| TonB-dependent receptor SusC | susC | 9 | 14 |
| Major outer membrane protein P. IA | porA | 9 | 10 |
| Succinate dehydrogenase flavoprotein subunit | sdhA | 2 | 9 |
| Outer membrane protein W | ompW | 7 | 8 |
| Putative outer membrane protein | Putative Omp | 8 | 8 |
| Elongation factor Tu | tufA | 2 | 8 |
| Outer membrane porin protein | Porin | 7 | 7 |
| 47 kDa outer membrane protein | omp 47KDa | 2 | 4 |
| Citrate synthase | gltA | 4 | 4 |
| Long-chain fatty acid transport protein | fadL | 3 | 4 |
| Outer membrane protein P1 | ompP1 | 4 | 4 |
| Elongation factor G | fusA | 3 | 4 |
| ATP synthase subunit b | atpF | 3 | 3 |
| DNA-directed RNA polymerase subunit beta | rpoB | 1 | 3 |
| Glycerol kinase | glpK | 3 | 3 |
| Outer membrane protein 40 | omp40 | 3 | 3 |
| Phosphoenolpyruvate carboxykinase [GTP] | pckG | 3 | 3 |
| Porin D | Porin D | 2 | 3 |
| Porin Omp2b | Porin Omp2b | 3 | 3 |
| Succinate--CoA ligase [ADP-forming] subunit beta | SUCLA2 | 3 | 3 |
| 2-oxoglutarate carboxylase large subunit | cfiA | 1 | 2 |
| 30S ribosomal protein S1 | rpsA | 1 | 2 |
| 3-methylmercaptopropionyl-CoA dehydrogenase | dmdC | 2 | 2 |
| 50S ribosomal protein L1 | rplA | 0 | 2 |
| 50S ribosomal protein L5 | rplE | 1 | 2 |
| Acetyl-coenzyme A synthetase | acs | 2 | 2 |
| ATP synthase subunit alpha | atpA | 0 | 2 |
| ATP synthase subunit beta | atpF | 1 | 2 |
| ATP synthase subunit beta 1 | atpD | 1 | 2 |
| Biopolymer transport protein ExbB | exbB | 2 | 2 |
| DNA-binding protein HU-beta | hupB | 2 | 2 |
| Flagellin | fliC | 2 | 2 |
| Fumarate reductase flavoprotein subunit | frdA | 1 | 2 |
| Ketol-acid reductoisomerase (NADP (+)) | ilvC | 2 | 2 |
| Major outer membrane prolipoprotein Lpp | lpp | 1 | 2 |
| Major outer membrane protein P. IB | porB | 2 | 2 |
| Malate dehydrogenase | MDH | 2 | 2 |
| Outer membrane protein | omp | 2 | 2 |
| Outer membrane protein A | ompA | 2 | 2 |

| Protein name | Gene name | FDR 1% | FDR 5% |
|---|---|---|---|
| Outer membrane protein IIIA | ropA | 2 | 2 |
| Outer membrane protein Omp38 | omp38 | 2 | 2 |
| Particulate methane monooxygenase alpha subunit | pmoB1 | 2 | 2 |
| Peroxiredoxin | Peroxiredoxi n | 1 | 2 |
| Phosphate-binding protein PstS | PstS | 2 | 2 |
| Porin | Porin | 0 | 2 |
| Protein oar | oar | 2 | 2 |
| Succinate--CoA ligase [ADP-forming] subunit alpha | sucD | 2 | 2 |
| Transcription termination/antitermination protein NusA | nusA | 0 | 2 |
| 60 kDa chaperonin | groL1 | 0 | 2 |
| V-type ATP synthase subunit C | atpC | 2 | 2 |
| 30S ribosomal protein S16 | rpsP | 1 | 1 |
| 30S ribosomal protein S3 | rpsC | 1 | 1 |
| 30S ribosomal protein S5 | rpsE | 1 | 1 |
| 30S ribosomal protein S7 | rpsG | 1 | 1 |
| 3-isopropylmalate dehydratase large subunit | IIL1 | 1 | 1 |
| 50S ribosomal protein L13 | rplM | 1 | 1 |
| 50S ribosomal protein L28 | rpmB | 1 | 1 |
| 5-methyltetrahydrofolate: corrinoid/iron-sulfur protein co-methyltransferase | acsE | 1 | 1 |
| Aconitate hydratase B | acnB | 1 | 1 |
| Adenylylsulfate reductase subunit alpha | aprA | 1 | 1 |
| Aerobic glycerol-3-phosphate dehydrogenase | GlpD | 1 | 1 |
| ATP synthase subunit c | atpC | 1 | 1 |
| ATP-dependent RecD-like DNA helicase | recD2 | 0 | 1 |
| Biotin transporter BioY | bioY | 0 | 1 |
| Calcium dodecin | Calcium dodecin | 1 | 1 |
| Carbon monoxide dehydrogenase/acetyl-CoA synthase subunit alpha | CODH/acs | 1 | 1 |
| Cation/acetate symporter ActP | actP | 1 | 1 |
| Chaperone protein DnaK | dnaK | 0 | 1 |
| Corrinoid/iron-sulfur protein large subunit | acsC | 1 | 1 |
| Cytochrome c-552 | cyt-c552 | 0 | 1 |
| DNA-binding protein HRm | HRm | 1 | 1 |
| Electron transfer flavoprotein subunit alpha | etfA | 1 | 1 |
| Electron transfer flavoprotein subunit beta | etfB | 1 | 1 |
| Enolase | eno | 0 | 1 |
| Ethanolamine ammonia-lyase heavy chain | eutB | 1 | 1 |
| Fatty acid oxidation complex subunit alpha | fadB | 1 | 1 |
| Fimbrial protein | fimA | 1 | 1 |
| GDP-6-deoxy-D-mannose reductase | rmd | 0 | 1 |
| Glutamyl-tRNA reductase | hemA | 0 | 1 |
| Glyceraldehyde-3-phosphate dehydrogenase 1 | GAPDH | 1 | 1 |

| Protein name | Gene name | FDR 1% | FDR 5% |
|---|---|---|---|
| GTP-binding protein TypA/BipA | TypA/BipA | 1 | 1 |
| Hydrogenase-1 large chain | hyaB | 0 | 1 |
| Inositol 2-dehydrogenase/D-chiro-inositol 3-dehydrogenase | iolG | 1 | 1 |
| Isocitrate dehydrogenase [NADP] | IDH1 | 1 | 1 |
| Isocitrate lyase | icl | 1 | 1 |
| Macrolide export protein MacA | macA | 0 | 1 |
| Maltoporin | lamB | 1 | 1 |
| Methylmalonyl-CoA mutase | mcm | 1 | 1 |
| Multidrug efflux pump subunit AcrA | acrA | 0 | 1 |
| NAD(P)H-quinone oxidoreductase subunit I chloroplastic | ndhI | 1 | 1 |
| NADP-dependent malic enzyme | maeB | 1 | 1 |
| Nitric oxide reductase subunit C | norC | 0 | 1 |
| Nitrogen regulatory protein | glnB | 1 | 1 |
| Nucleoside diphosphate kinase | ndk | 0 | 1 |
| Oligopeptide-binding protein AppA | appA | 0 | 1 |
| Outer membrane protein 41 | omp41 | 1 | 1 |
| Outer membrane protein C | ompC | 1 | 1 |
| Outer membrane protein P6 | ompP6 | 1 | 1 |
| Outer membrane protein PagN | pagN | 1 | 1 |
| Outer membrane protein X | ompX | 1 | 1 |
| 5,10-methylenetetrahydromethanopterin reductase | mer | 1 | 1 |
| Putative adenylyl-sulfate kinase | cysC | 0 | 1 |
| Putative glutamine ABC transporter permease protein GlnM | GlnM | 0 | 1 |
| Putative phospholipase A1 | pldA | 0 | 1 |
| Pyruvate dehydrogenase E1 component | PDHA1 | 1 | 1 |
| Ribonuclease HII | rnhB | 0 | 1 |
| Ribulokinase | araB | 0 | 1 |
| RNA polymerase sigma factor RpoD | rpoD | 1 | 1 |
| S-layer protein SlpA | slpA | 1 | 1 |
| Superoxide dismutase [Fe] | SODB | 1 | 1 |
| Thioredoxin | Thioredoxin | 1 | 1 |
| Transcription-repair-coupling factor | mfd | 0 | 1 |
| Trigger factor | tig | 1 | 1 |
| Tubulin-like protein CetZ | cetZ | 1 | 1 |
| V-type ATP synthase alpha chain | atpA | 1 | 1 |

### Table S11

*Table 11. List of all genera associated with three or less expressed proteins.*

| Class | Genus | Number of expressed proteins |
|---|---|---|
| Acidobacteria | Candidatus Solibacter | 3 |
| Actinobacteria | Aurantimicrobium | 3 |
| | Ilumatobacter | 3 |
| | Nocardiopsis | 3 |
| | Tessaracoccus | 3 |
| Alphaproteobacteria | Agrobacterium | 3 |
| | Defluviicoccus | 3 |
| | Georhizobium | 2 |
| | Caulobacter | 2 |
| | Croceicoccus | 2 |
| | Paracoccus | 2 |
| | Rhodopseudomonas | 2 |
| | Roseomonas | 2 |
| | Shinella | 2 |
| | Stella | 2 |
| | Tabrizicola | 2 |
| Bacteroidetes | Bacteroidales bacterium CF | 2 |
| | Cloacibacterium | 2 |
| | Lacinutrix | 2 |
| | Alistipes | 2 |
| | Dysgonomonas | 2 |
| | Elizabethkingia | 2 |
| | Filimonas | 2 |
| | Flavobacteriaceae bacterium UJ101 | 2 |
| | Flavobacterium | 2 |
| | Labilibaculum | 2 |
| | Lutibacter | 2 |
| | Parabacteroides | 2 |
| | Petrimonas | 2 |
| | Prevotella | 2 |
| | Rhodothermaceae bacterium RA | 2 |
| | Salinivirga | 2 |
| | Sphingobacterium | 1 |
| Betaproteobacteria | Comamonas | 3 |
| | Ephemeroptericola | 3 |
| | beta proteobacterium CB | 2 |
| | Delftia | 1 |
| | Polynucleobacter | 1 |
| | Ramlibacter | 1 |
| | Rhodoferax | 1 |
| | Serpentinomonas | 1 |

| Class | Genus | Number of expressed proteins |
| --- | --- | --- |
| | Sulfurimicrobium | 1 |
| | Verminephrobacter | 1 |
| | Achromobacter | 1 |
| | Cupriavidus | 1 |
| | Ferriphaselus | 1 |
| | Iodobacter | 1 |
| | Methylibium | 1 |
| | Methyloversatilis | 1 |
| | Nitrosomonas | 1 |
| | Pigmentiphaga | 1 |
| | Sulfuricella | 1 |
| | Sulfuriferula | 1 |
| | Undibacterium | 1 |
| Chlamydiae | Neochlamydia | 1 |
| Chlorobi | Ignavibacterium | 1 |
| | Prosthecochloris | 1 |
| Chloroflexi | Pelolinea | 1 |
| Deltaproteobacteria | Desulfosarcina | 1 |
| | Desulfuromonas | 1 |
| | Desulfobulbus | 1 |
| | Desulfococcus | 1 |
| | Anaeromyxobacter | 1 |
| | Desulfobacterium | 1 |
| | Desulfomonile | 1 |
| | Geobacter | 1 |
| | Haliangium | 1 |
| | Sorangium | 1 |
| Epsilonproteobacteria | Sulfuricurvum | 1 |
| | Sulfurimonas | 1 |
| | Pseudoarcobacter | 1 |
| | Sulfurospirillum | 1 |
| | Sulfurovum | 1 |
| Firmicutes - Bacilli | Thermobacillus | 1 |
| Firmicutes - Clostridia | Caldanaerobacter | 1 |
| | Caloramator | 1 |
| | Caproiciproducens | 1 |
| | Moorella | 1 |
| | Syntrophomonas | 1 |
| | Thermincola | 1 |
| Fusobacteria | Ilyobacter | 1 |
| Gammaproteobacteria - Enterobacteria | Escherichia | 1 |
| | Shigella | 1 |
| | Shimwellia | 1 |

| Class | Genus | Number of expressed proteins |
| --- | --- | --- |
| Gammaproteobacteria - Others | Acinetobacter | 1 |
| | Aeromonas | 1 |
| | Azotobacter | 1 |
| | Methylomonas | 1 |
| | Pseudomonas | 1 |
| | Aquicella | 1 |
| | Dokdonella | 1 |
| | Dyella | 1 |
| | Entomomonas | 1 |
| | Methylocaldum | 1 |
| | Methylomicrobium | 1 |
| | Microbulbifer | 1 |
| | Oblitimonas | 1 |
| | Permianibacter | 1 |
| | Saccharophagus | 1 |
| | Tatlockia | 1 |
| | Thermomonas | 1 |
| | Thioflavicoccus | 1 |
| | Xanthomonas | 1 |
| Lentisphaerae | Victivallales bacterium CCUG 44730 | 1 |
| Saccharibacteria | Candidatus Saccharibacteria oral taxon TM7x | 1 |
| Spirochaetes | Salinispira | 1 |
| | Treponema | 1 |
| | Turneriella | 1 |
| Synergistetes | Cloacibacillus | 1 |
| Unclassified Bacteria | Candidatus Campbellbacteria bacterium GW2011_OD1_34_28 | 1 |

### Table S12

*Table 12. List of associated expressed proteins to Paucimonas.*

| Gene | Number | Description | KO number |
|---|---|---|---|
| atpD | 1 | ATP synthase subunit beta | K02112 |
| cfiA | 1 | 2-oxoglutarate carboxylase large subunit | K01960 |
| cysC | 1 | putative adenylyl-sulfate kinase | K00955 |
| etfB | 1 | Electron transfer flavoprotein subunit beta | K03521 |
| fadB | 1 | Fatty acid oxidation complex subunit alpha | K01825 |
| GAPDH | 1 | Glyceraldehyde-3-phosphate dehydrogenase 1 | K00134 |
| GlpD | 1 | Aerobic glycerol-3-phosphate dehydrogenase | K00111 |
| HRm | 1 | DNA-binding protein HRm | K03530 |
| maeB | 1 | NADP-dependent malic enzyme | K00029 |
| ompP6 | 1 | Outer membrane protein P6 | K03640 |
| porin D | 1 | Porin D | K18093 |
| PstS | 1 | Phosphate-binding protein PstS | K02040 |
| rplE | 1 | 50S ribosomal protein L5 | K02931 |
| rplM | 1 | 50S ribosomal protein L13 | K02871 |
| rpmB | 1 | 50S ribosomal protein L28 | K02902 |
| rpsA | 2 | 30S ribosomal protein S1 | K02945 |
| rpsC | 1 | 30S ribosomal protein S3 | K02982 |
| rpsE | 1 | 30S ribosomal protein S5 | K02988 |
| rpsP | 1 | 30S ribosomal protein S16 | K02959 |
| sucD | 1 | Succinate--CoA ligase [ADP-forming] subunit alpha | K01902 |
| SUCLA2 | 1 | Succinate--CoA ligase [ADP-forming] subunit beta | K01903 |
| tig | 1 | Trigger factor | K03545 |
| tufA | 2 | Elongation factor Tu | K02358 |
| TypA/BipA | 1 | GTP-binding protein TypA/BipA | K06207 |

### Figure S5

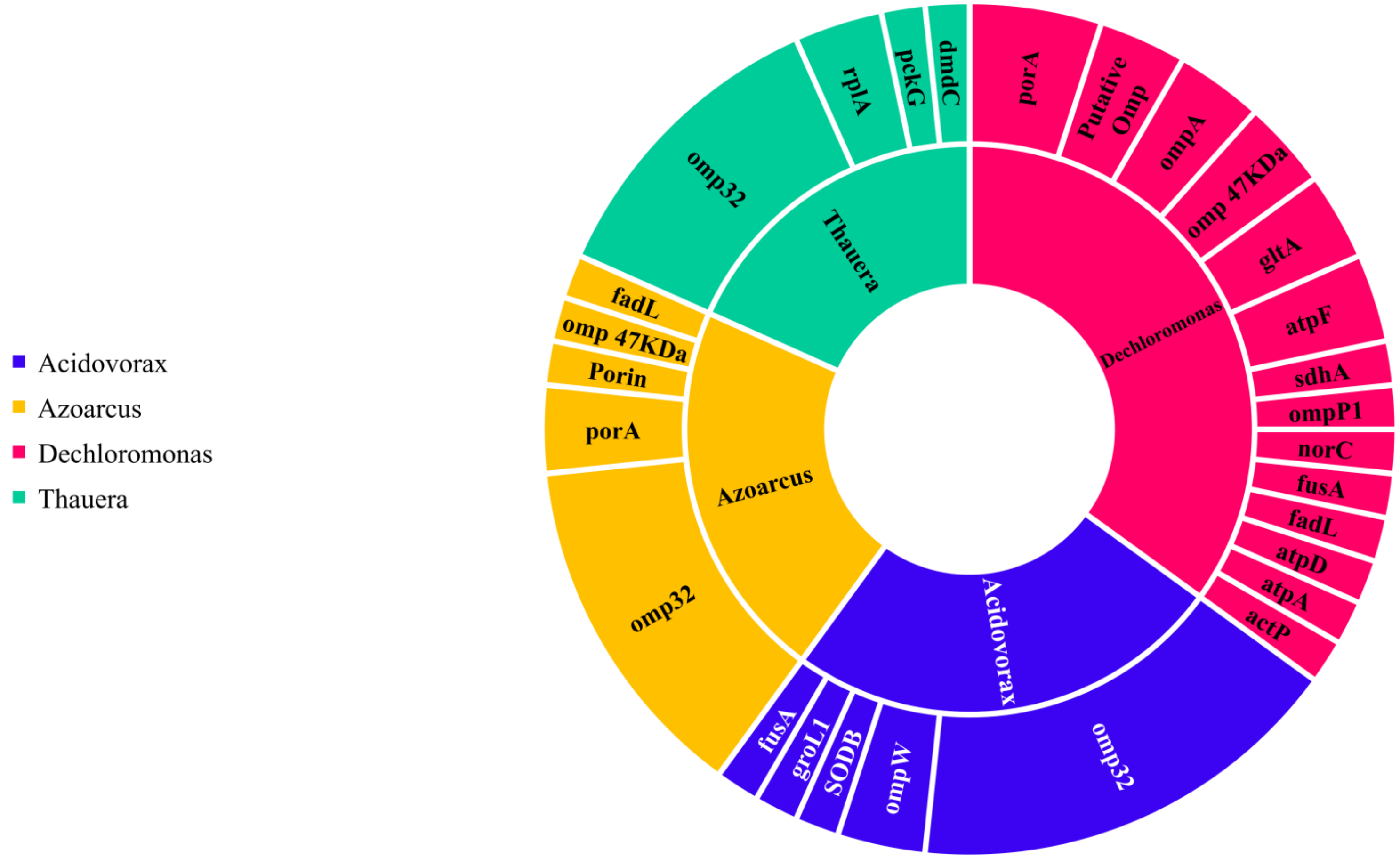


*Figure 5. Related expressed genes for top-ranked genera identified by proteomics, actP=Cation/acetate symporter, atpA=ATP synthase subunit alpha, atpD=ATP synthase subunit beta 1, atpF= ATP synthase subunit b, dmdC=3-methylmercaptopropionyl-CoA dehydrogenase, fadL= Long-chain fatty acid transporter, fusA=Elongation factor G, gltA=Citrate synthase, groL1= 60 kDa chaperonin, norC=Nitric oxide reductase subunit C, omp 47KDa= 47 kDa outer membrane protein, omp32=Outer membrane porin protein 32, ompA= Outer membrane protein A, ompP1= Outer membrane protein P1, omp W=Outer membrane protein W, pckG=Phosphoenolpyruvate carboxykinase [GTP], porA=Major outer membrane protein P.IA, Porin=Outer membrane porin protein, Putative Omp=Putative outer membrane protein, rplA=50S ribosomal protein L1, sdhA=Succinate dehydrogenase flavoprotein subunit, SODB=Superoxide dismutase [Fe]*
