## Supplementary-File 2 for "Looking for lipases and lipolytic organisms in low-temperature anaerobic reactors treating domestic wastewater"

### Protein downstream processes

#### VSS measurement

Microfiber Whatman filter papers were first dried in an oven at 105 °C for 15 min then in a furnace at 550 °C for 5 min. they were later cooled down in a desiccator, labeled by a soft pencil and weighed to constant value. 10 ml and 1 ml of bulk liquid and biofilm from AnMBRs were filtered, respectively and first dried for 1 hr at 105 °C then at 550 °C for 5 min. After the ignition, filter papers were cooled down in a desiccator and weighed to the constant value. The initial weight of the empty filter papers was subtracted from the weight obtained after the ignition and reported as g/l.

#### Protein extraction

From each reactor, 10 ml of bulk liquid and 1 ml of biofilm were collected and transferred to individual 50 ml conical centrifuge tubes. 9 ml autoclaved distilled water was added to biofilm-containing tubes to retain the same volume. 5 gr cation exchange resin (DOWEX, 50X8, 20-50 mesh, $Na^+$ form, strong acidic, Sigma Aldrich) pre-washed for 1 h in sample buffer (2 mM $Na_3PO_4$, 4 mM $NaH_2PO_4$, 9 mM NaCl and 1 mM KCl at pH=7) along with 10 µl Triton X-100 (final concentration of 0.1% v/v) was added to each tube. Samples were shaken for 1.5 h at 400 rpm and 4°C and then centrifuged twice at the same temperature (20 min at 15,000g and 10 min at 10,000g). The supernatant was collected for protein quantification and precipitation.

#### Protein quantification

Pierce™ Modified Lowry Protein Assay Kit was used for measuring the concentration of proteins in supernatant and plotting the standard curve of Bovine serum albumin (BSA) (Figure

1). The BSA with concentration of 2 mg/ml was diluted into various ranges of 1, 5, 25, 125, 250, 500, 750, 1000 and 1500 µg/ml according to the kit instructions. 0.2 ml of each dilution was mixed with 1 ml of modified Lowry reagent, vortexed and incubated for 10 min at room temperature. Finally, 0.1 ml of phenol reagent (already diluted with distilled water to yield 1 N solution) was added to each sample, vortexed and incubated at room temperature for another 30 min. The absorbances were read at 750 nm and plotted against concentrations of diluted samples for further calculations.

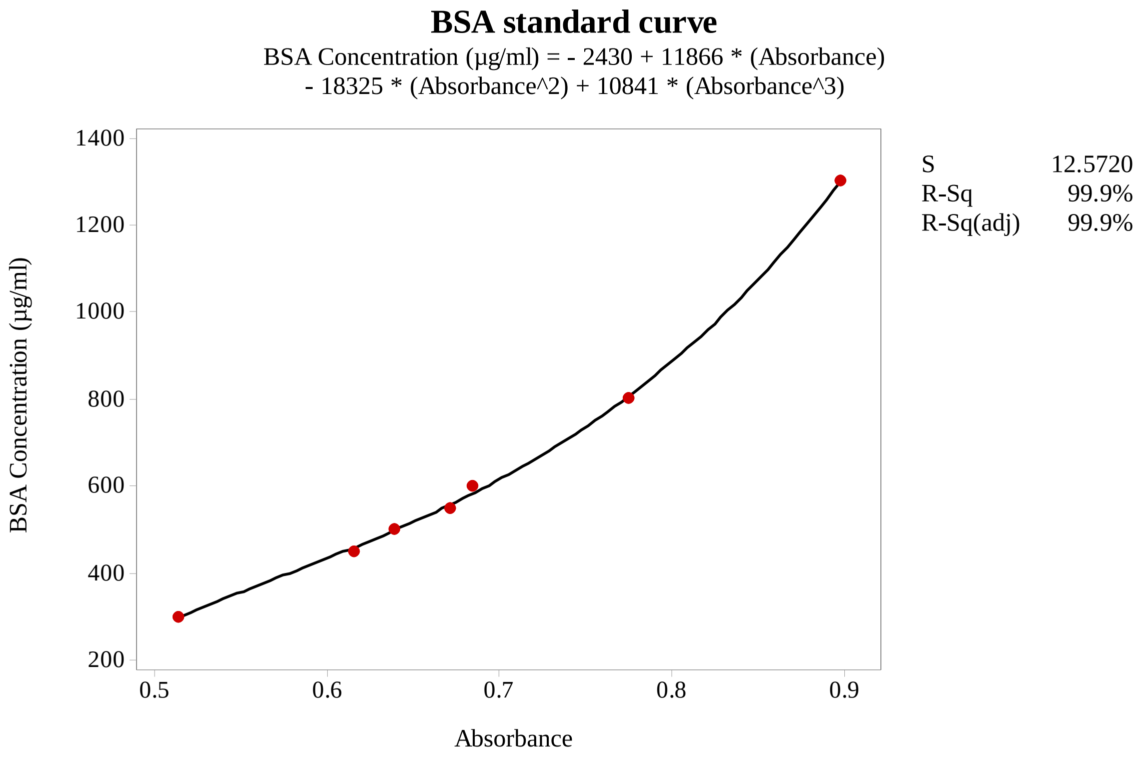


*Figure 1. BSA standard curve, cubic regression model, Minitab 18.*

### Protein precipitation

1 part of supernatant was mixed by 4 part of ice-cold methanol and vortexed. 1 part of ice-cold chloroform and 3 parts of cold distilled water were added respectively and vortexed too. The mixture was then centrifuged for 1 min at 15500g and 4 ºC to form three phases (proteins form a circular flake in the interface of water and chloroform). The top aqueous layer containing salts and hydrophilic contaminants was carefully removed by pipette. 4 part of methanol was

added again and after being vortexed, the mixture was centrifuged for 5 min at 15500g and 4 ºC. After removing the supernatant, the pellets were air-dried and stored at -80 °C for further analysis.

**1D SDS-PAGE**

100 µl of BME and 900 µl of Laemmli buffer were mixed and 10 µl of the mixture was added to each tube containing protein pellets (defrosted in room temperature). The tubes were sonicated for 20 min at cool temperature, then heated at 60 °C for 5 min and centrifuged for 10 min at 4 °C. 10 µl of supernatant was injected into wells (4–15% Mini-PROTEAN® TGX™ Precast Protein Gels, 15-well, 15 µl) and run for 5 min at 120 V (Bio-Rad Mini-PROTEAN®). The gel was removed from the tank, was immersed in distilled water, microwaved for 1 min, and shaken at 360 rpm for 1 min (PMS-1000i Microplate Shaker, Grant Instruments™). The water was removed, and the washing/microwaving/shaking procedure was repeated for three times. After removing the water, the gel was stained by 60 ml Bio-Safe Coomassie Brilliant Blue G-250 (microwaved for 1 min and shaken for 5 min at 360 rpm) and destained overnight in distilled water at 360 rpm and room temperature. Destained gel was stored at 4 °C in 20 mM NaCl solution before in-gel digestion.

**In-gel digestion**

Each 1D SDS-PAGE band was excised with a clean scalpel, diced into 1x1x1 mm cubes, and transferred to a clean microcentrifuge tube. Gel pieces were destained by mixture of 50mM ammonium bicarbonate and acetonitrile (50%). The destained buffer was removed and exchanged until the gel pieces were clear. As a digest control, a molecular weight marker band was also excised. Proteins were reduced with 10 mM dithiothreitol for 30 min at 60˚C to break disulphide bridges. This was followed by alkylation with 50 mM iodoacetamide for 30 min at room temperature in the dark to prevent disulphide reformation. Gel pieces were washed in 50mM ammonium bicarbonate and then dehydrated with 3 washes of 100 μL of acetonitrile. Residual moisture was removed from gel pieces in a vacuum drier. Proteins were digested by the addition of trypsin added at a ratio of 30:1 (protein: trypsin), buffered with 50 mM ammonium bicarbonate and incubated for 16 hours at 37 ˚C. The digest was stopped by the addition of 10% Trifluoroacetic acid (TFA) to a final concentration of 0.5%, shaken for 30 mins, 750 rpm. The liquid containing hydrophilic peptides was transferred to a fresh microcentrifuge tube. 80% acetonitrile with 2% TFA was then added to the gel pieces and shaken for 30 min at 750 rpm. This dehydrates the gel pieces and removes hydrophobic peptides from the gel. The solution containing hydrophobic peptides was pooled with the hydrophilic peptide mix. The peptide solution was dried in a centrifugal evaporator, peptides were dissolved in 3% acetonitrile, and 0.1% TFA. The resulting peptide solutions were desalted using home packed C18 stage tips (Rappsilber et al. 2007). The sample was dissolved in 50 μL of 3% acetonitrile, 0.1% TFA giving the final concentration of ~1μg/μL.

**Nano LC-MS/MS**

About 1 µg of a protein digest was loaded onto a UltiMate 3000 RSLC nano HPLC and peptides separated with a 97 min nonlinear gradient (3-40%, 0.1% formic acid). Samples were first loaded onto a 300 µm x 5mm C18 PepMap C18 trap cartridge in 0.1% formic acid at 25 µl/min and passed on to an in-house made 75 µm x 15cm C18 column (ReproSil-Pur Basic-C18-HD, 3 µm, Dr. Maisch GmbH) at 400nl/min. The eluent was directed to an Ab-Sciex TripleTOF 6600 mass spectrometer through the AB-Sciex Nano-Spray 3 source, fitted with a New Objective FS360-20-10 emitter. For data-dependent data acquisition (DDA), MS1 data was acquired within a range of 400-1250m/z (250 ms accumulation time), followed by MS2 of Top 30 precursors with charge states between 2 and 5 (total cycle time 1.8s). Product ion spectra (50 ms accumulation time) were acquired within a range of 100-1500m/z, using rolling collision energy for precursors which exceed 150 cps. Precursor ions were excluded for 15s after one occurrence. The acquired DDA data was searched against the metagenomics sequence database.
